# CHD7 binds to insulators during neuronal differentiation

**DOI:** 10.1101/2025.03.28.646031

**Authors:** Jingyun Qiu, Azadeh Jadali, Edward Martinez, Zhichao Song, Julie Z. Ni, Kelvin Y. Kwan

**Affiliations:** Department of Cell Biology & Neuroscience, Rutgers University, Piscataway, NJ 08854, USA; Stem Cell Research Center and Keck Center for Collaborative Neuroscience, Rutgers University, Piscataway, NJ 08854, USA; Sampled, Piscataway, NJ 08854, USA; Department of Molecular Biology and Biochemistry, Rutgers University, Piscataway, NJ 08854, USA

**Author notes:** These authors contributed equally (Jingyun Qiu, Azadeh Jadali). Corresponding author Address: 604 Allison Rd., Nelson Labs D250, Piscataway NJ 08854.

## Abstract

Spiral ganglion neurons (SGNs) are crucial for hearing, and the loss of SGNs causes hearing loss. Stem cell-based therapies offer a promising approach for SGN regeneration and require understanding the mechanisms governing SGN differentiation. We investigated the chromatin remodeler CHD7 in neuronal differentiation using immortalized multipotent otic progenitor (iMOP) cells. We demonstrated that CHD7 knockdown impaired neuronal differentiation. Genome-wide analysis revealed CHD7 binding at diverse *cis*-regulatory elements, with notable enrichment at sites marked by the insulator-binding protein CTCF between topologically associating domains (TADs). Insulators marked by the enrichment of CHD7 and CTCF resided near genes critical for neuronal differentiation, including *Mir9-2*. Targeting these regulatory regions in iMOPs with CRISPR interference (CRISPRi) and activation (CRISPRa) increased miR-9 transcription, irrespective of the method. Blocking the CHD7 and CTCF marked sites suggested that the elements function as insulators to regulate gene expression. The study highlights CHD7 activity at insulators and underscores an unreported mechanism for promoting neuronal differentiation.

## INTRODUCTION

Exposure to loud sounds results in the gradual loss of spiral ganglion neurons (SGNs) (Kujawa and Liberman 2006, Kujawa and Liberman 2009). Once lost, SGNs cannot regenerate. The loss of SGNs impairs auditory function and results in hearing loss (Shi and Edge 2013). No treatments are currently available for SGN loss, but stem cell replacement therapy is a strategy to alleviate auditory neuropathy (Corrales, Pan et al. 2006, Chen, Jongkamonwiwat et al. 2012). After the chemical ablation of SGNs, the engraftment of otic neural progenitors derived from human pluripotent stem cells into the inner ear allows for partial hearing recovery (Chen, Jongkamonwiwat et al. 2012). Robust functional hearing recovery requires nascent SGNs to attain the proper morphology, axonal targeting, and synaptic specificity. Few engrafted stem cell-derived neurons display these properties; some may develop into inappropriate cell types or teratomas that reduce the therapeutic efficacy of stem cell replacement (Nishimura, Nakagawa et al. 2012). Identifying the molecular mechanisms that facilitate the differentiation of otic progenitors into SGNs will accelerate efforts for stem cell or progenitor cell-based therapies for regeneration.

Cell fate transitions are orchestrated by changes in gene expression, which in turn is governed by *cis*-regulatory elements such as promoters, enhancers, silencers, and insulators. The activities of these elements are intricately regulated by the epigenomic landscape, including histone modifications and chromatin accessibility (Strahl and Allis 2000, Kouzarides 2007, Li, Carey et al. 2007). For instance, histone acetylation plays a crucial role in modulating the activity of distal regulatory regions, influencing the appropriate enhancer repertoire during cell fate specification (Zaghi, Banfi et al. 2023). Chromatin remodeling proteins, like CHD8, can interact with insulator-binding proteins such as CTCF to modulate gene expression by affecting the function of *cis*-regulatory elements (Ishihara, Oshimura et al. 2006). Given the importance of chromatin dynamics in cell fate determination, we speculate that the chromatin remodeler CHD7 plays a critical role in SGN differentiation.

CHD7 is an ATP-dependent chromatin remodeling protein known to reposition nucleosomes (Bouazoune and Kingston 2012). CHD7 interacts with methylated histones via its chromodomain (Jacobs and Khorasanizadeh 2002, Nielsen, Nietlispach et al. 2002) and has been implicated in regulating *cis*-regulatory elements in various cell types. In human colorectal carcinoma cells, human neuroblastoma cells, and murine ES cells, CHD7 is enriched at cell-type specific enhancers (Schnetz, Bartels et al. 2009). In human neural crest cells, human induced pluripotent stem cells, and human neuroepithelial cells, CHD7 binding sites are marked by H3K4me3, H3K4m1, H3K27ac, and EP300 (p300) (Sanosaka, Okuno et al. 2022). Furthermore, CHD7 interacts with the insulator-associated protein CTCF through its BRK domain (Allen, Religa et al. 2007), suggesting a potential role with the DNA-binding protein. The association of CHD7 to different *cis*-regulatory elements - enhancers, promoters, and insulators - depends on the cellular context.

Pathogenic variants in the *CHD7* gene cause CHARGE syndrome with haploinsufficiency as the proposed underlying mechanism (Vissers, van Ravenswaaij et al. 2004). Among the developmental deficits observed in CHARGE syndrome, ear anomalies resulting in hearing loss are highly penetrant (Zentner, Layman et al. 2010). Sensorineural hearing loss in CHARGE syndrome is likely related to improper SGN development (Thelin, Mitchell et al. 1986, Thelin and Fussner 2005). Conditional knockout mouse studies implicate CHD7 in SGN production or differentiation (Hurd, Capers et al. 2007, Hurd, Poucher et al. 2010, Hurd, Adams et al. 2011) and protect SGNs from stress-induced degeneration (Ahmed, Moon et al. 2021). To investigate CHD7’s role in otic progenitor cell differentiation into neurons, we employed a clonally-derived immortalized multipotent otic progenitor (iMOP) cell line as a model system. This cellular model allows us to dissect the function of CHD7 and examine its impact on the molecular events underlying SGN differentiation.

## RESULTS

### CHD7 associates with SOX2 gene regulatory network in iMOPs

iMOP cells derived from SOX2-expressing primary progenitor cells of the developing inner ear have the capacity for self-renewal and can differentiate into otic cell types (Kwan, Shen et al. 2015). Transcriptome comparison of iMOP cells to SOX2-expressing primary otic progenitors obtained from embryonic cochlea showed high similarity (Kwan, Shen et al. 2015). iMOP cells retain the potential to differentiate into bipolar and pseudounipolar neurons that express markers for spiral ganglion neurons (SGNs) (Kwan, Shen et al. 2015, Jadali, Song et al. 2016). Proliferating iMOP cells and iMOP-derived neuron cultures were used for RNA-seq to investigate changes in the transcriptome during the neuronal differentiation of iMOPs. Proliferating iMOP cells grew as otospheres, and only a small population of cells expressed low levels of the cyclin-dependent kinase inhibitor 1B (CDKN1B) to transiently inhibit cell cycle progression. The vast majority of proliferating iMOP cells did not express neuronal β-tubulin (TUBB3), as evidenced by the lack of fluorescence signal in Hoechst-labeled cells (Fig. 1A). Only 0.83% of cells displayed detectable TUBB3. The data suggests that proliferating iMOP cells contain progenitor cells with a low tendency towards spontaneous differentiation. iMOP cells cultured under neuronal differentiation conditions expressed high levels of CDKN1B and displayed pseudounipolar and bipolar neurites marked by TUBB3 (Fig. 1B), consistent with cell cycle exit and expression of a neuronal marker. Over 98.1% of cells in differentiating cultures expressed both CDKN1B and TUBB3. From this point forward, we refer to proliferating iMOPs as progenitors and iMOP-derived neurons as neurons.

**Fig. 1.**
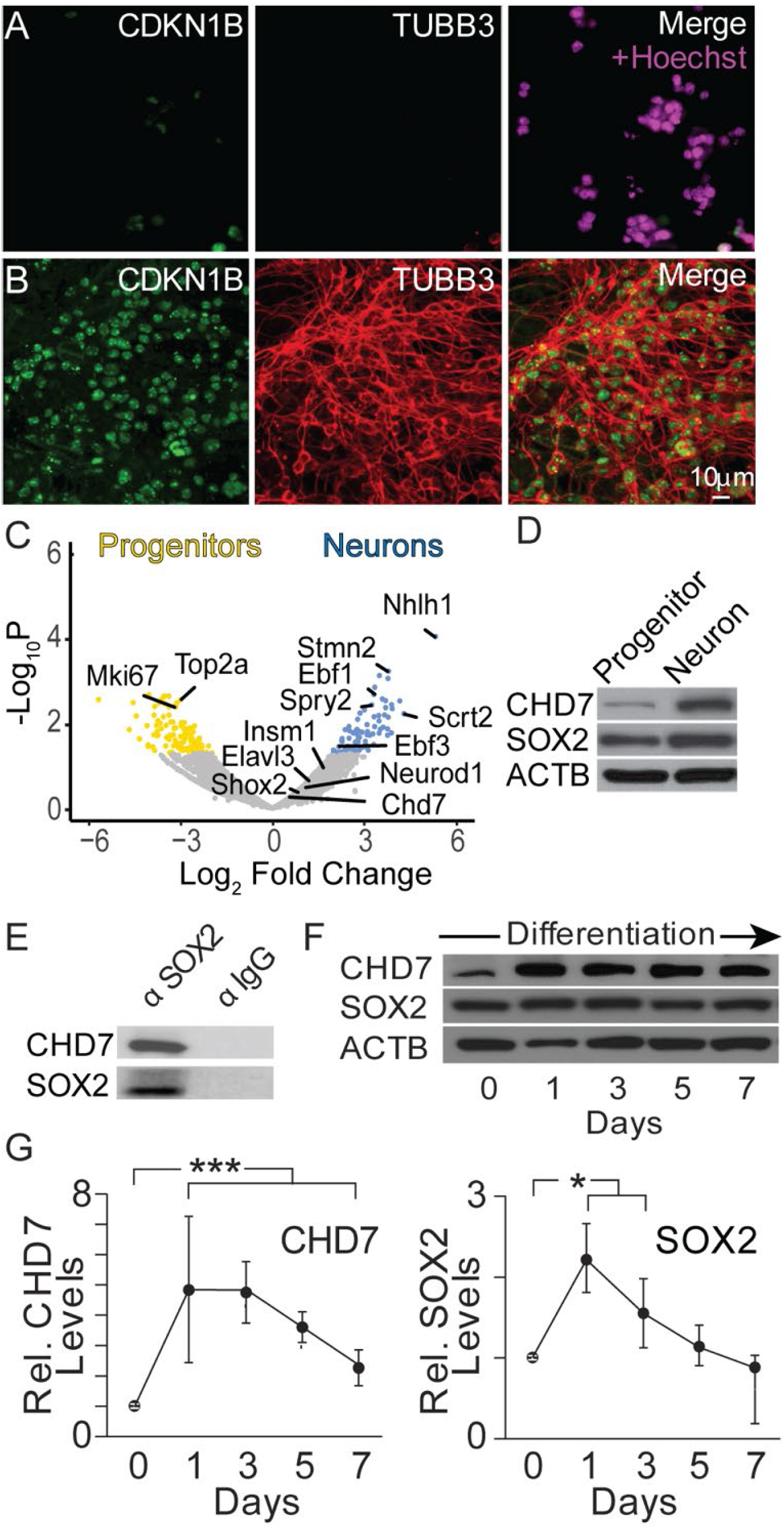
CHD7 and SOX2 expression in differentiating iMOPs. Fluorescent images of CDKN1B, TUBB3, and Hoechst-labeled nuclei in (A) proliferating iMOPs (progenitors) and (B) iMOP-derived neurons (neurons). (C) Bulk RNA-seq analysis comparing gene expression of progenitors and neurons. Genes associated with proliferation and neuronal differentiation were marked. CHD7, a chromatin remodeling protein, is present in both cell types. (D) Western blot of CHD7, SOX2, and ACTB using extracts from progenitors and neurons (n=3 independent experiments). (E) Western blot of CHD7 and SOX2 after immunoprecipitation with a SOX2 antibody or an IgG antibody (n=3 independent experiments). (F) Time course of CHD7, SOX2, and ACTB expression levels during neuronal differentiation of iMOP cells by Western blot (n=5 independent experiments). (G) CHD7 and SOX2 signals were quantified from Western blots at different time points during iMOP neuronal differentiation after normalizing to ACTB (n=5, independent experiments). Student’s T-test was used for statistical analysis, * p<0.05, *** p<0.001.

Using these cultures, progenitors and neurons were harvested, RNA-seq was performed, and differentially expressed genes from proliferating and differentiating iMOP cultures were identified (Fig. 1C). Sequence reads from individual transcripts were used to measure RNA levels. Transcripts with > 2-fold increase were highlighted. Statistically significant enriched genes (p<0.05) included *Nhlh1* and *Stmn2*, genes identified in early SGNs (Petitpre, Faure et al. 2022) and *Scrt2* (Li, Li et al. 2020), respectively, while progenitors showed robust expression of markers associated with proliferating cells such as *Mki67* (Gerdes, Schwab et al. 1983) and *Top2a* (Nitiss 1998). *Chd7* transcript was detected in both progenitors and neurons but showed no significant change. Gene ontology analysis using enriched genes in progenitors corresponded to biological processes involved in cell division, cell cycle, and chromatin organization. Genes enriched in neurons corresponded to sensory organ, axon, and neuron projection development (Fig. S1). RNA-seq analysis suggests that enriched genes in progenitors and neurons correspond to cellular processes indicative of their cell state.

CHD7 is a chromatin remodeling factor that regulates inner ear neurogenesis (Hurd, Poucher et al. 2010). CHD7 can bind to short linker DNA with high affinity and slide along the nucleosome, allowing it to reposition nucleosomes (Manning and Yusufzai 2017). Aside from binding to linker DNA, CHD7 associates with transcription factors that bind to specific regulatory regions. During cochlear development, the SOX2 and CHD7 gene regulatory network was recently established *in vivo* (Gao, Skidmore et al. 2024). To determine if CHD7 and SOX2 proteins are expressed in neurons, Western blots were performed on lysates obtained from progenitors and neurons for CHD7, SOX2, and ACTB (β-Actin). CHD7 protein levels increased in differentiating iMOP cultures relative to proliferating iMOP cultures (Fig. 1D). In neural stem cells, SOX2 and CHD7 were shown to interact (Engelen, Akinci et al. 2011). Immunoprecipitation with a SOX2 antibody followed by Western blotting was performed using neuronal lysates to determine if CHD7 and SOX2 biochemically interact in iMOPs similar to neural stem cells and developing otic cells. Immunoprecipitates were probed with CHD7 antibodies to assay for biochemical interaction between SOX2 and CHD7 in iMOP cells. Western blot analysis demonstrated the association of CHD7 with SOX2 in the immunoprecipitate. As a negative control, a secondary antibody was used for immunoprecipitation (Fig. 1E). We showed that CHD7 associates with the transcription factor SOX2 in iMOPs, supporting the *in vivo* studies.

CHD7 may be dynamically expressed during neuronal differentiation and provides insight into when it functions. To determine the temporal expression of CHD7 in differentiating iMOP cultures, cell lysates from cultures at different time points after induction of neuronal differentiation were collected. Western blot detected CHD7 and SOX2 during the 7-day differentiation period. Signal intensities were normalized to ACTB (Fig. 1F). Compared to the initial day of differentiation (Day 0), CHD7 showed a 4.8- and 4.7-fold enrichment on day 1 and day 3 after differentiation, respectively, then decreased to 3.6- and 2.3-fold enrichment on day 5 and day 7. A significant increase in CHD7 levels was observed for all time points relative to the onset of neuronal differentiation (p<1x10^-3^) (Fig. 1G). A similar expression profile was observed for SOX2. These results suggest that CHD7 and SOX2 are upregulated and could biochemically function together during neuronal differentiation. Despite not observing changes in the *Chd7* transcript levels, our data shows that CHD7 is dynamically expressed and suggests post-transcriptional control of CHD7 expression. These data also show that a key gene regulatory network – the CHD7-SOX2 axis - is present in iMOP during neuronal differentiation.

### CHD7 is required for neuronal differentiation of iMOP cells

To test whether CHD7 is essential for neuronal differentiation in iMOPs, we wanted to mimic CHD7 haploinsufficiency in CHARGE syndrome during neuronal differentiation using a knockdown strategy. Three unique *Chd7* shRNAs (*Chd7* shRNA1-3), including a previously published shRNA (Engelen, Akinci et al. 2011) and a scrambled shRNA, were inserted into a lentiviral vector containing a blasticidin resistance gene (pLKO.1-blast). Scrambled and *Chd7* shRNA oligos were cloned into pLKO.1-blast. Differentiating iMOP cells were transduced with individual lentivirus, selected in blasticidin, and total RNA harvested from cultures for qPCR. *Chd7* transcript levels were normalized to *Gapdh* levels. Normalized transcript levels from scrambled shRNA and *Chd7* shRNA transduced cultures were compared. *Chd7* shRNA1-3 was individually used to transduce cultures. *Chd7* transcript levels from the *Chd7* shRNA1-3 transduced cultures showed 0.13-, 0.21-, and 0.15-fold reduction relative to scrambled shRNA (Fig. S2A). The *Chd7* shRNA1 knockdown described in a previous report showed the most robust knockdown (Engelen, Akinci et al. 2011). *Chd7* shRNA1 was subsequently used to perform experiments. Western blot analysis showed decreased CHD7 protein levels after knockdown. After selecting infected cells, lysates were harvested from scrambled shRNA, or *Chd7* shRNA lentivirus transduced neurons for Western blot (Fig. S2B). Western blot signals from cell lysates showed that CHD7 protein levels were reduced to 0.32-fold in *Chd7* shRNA transduced cells compared to scrambled shRNA transduced cells (Fig. S2C). Primers for qPCR and shRNA sequences for *Chd7* knockdown are provided (Fig. S2D, Fig. S2E). These results show that CHD7 protein levels diminished after introducing *Chd7* shRNA.

Immunostaining was performed using antibodies against CHD7 to test whether CHD7 was present in progenitors and neurons. In progenitors, nuclear CHD7 labeling was observed. At the same time, phalloidin was used as a counterstain to label actin filaments in the cytoplasm (Fig. 2A). In neurons, CHD7 marked the nucleus of TUBB3 labeled cells (Fig. 2B). To determine whether acute *Chd7* knockdown affected iMOP differentiation, *Chd7* shRNA or scrambled shRNA was inserted after the U6 promoter in a GFP-expressing plasmid (pLKO.3G). Scrambled and *Chd7* shRNA oligos were used for cloning into pLKO.3G (Fig. S2E). The plasmids were transfected into iMOP cells 24 hours after initiating neuronal differentiation. GFP fluorescence highlighted transfected cells, marked cell bodies and neurites, and allowed sparse labeling of cells for analysis. After transfection, the iMOP-derived neuronal cultures were subjected to TUBB3 labeling. In scrambled shRNA cultures, both transfected and non-transfected cells displayed long TUBB3 neuronal processes (Fig. 2C). In contrast, *Chd7* shRNA transfected cells labeled with GFP expressed low levels of TUBB3 while untransfected cells that served as an internal control displayed well-defined TUBB3+ processes (Fig. 2D). These data indicate that CHD7 is present in both progenitor and neuronal cells and that its depletion impaired neuronal differentiation.

**Fig. 2.**
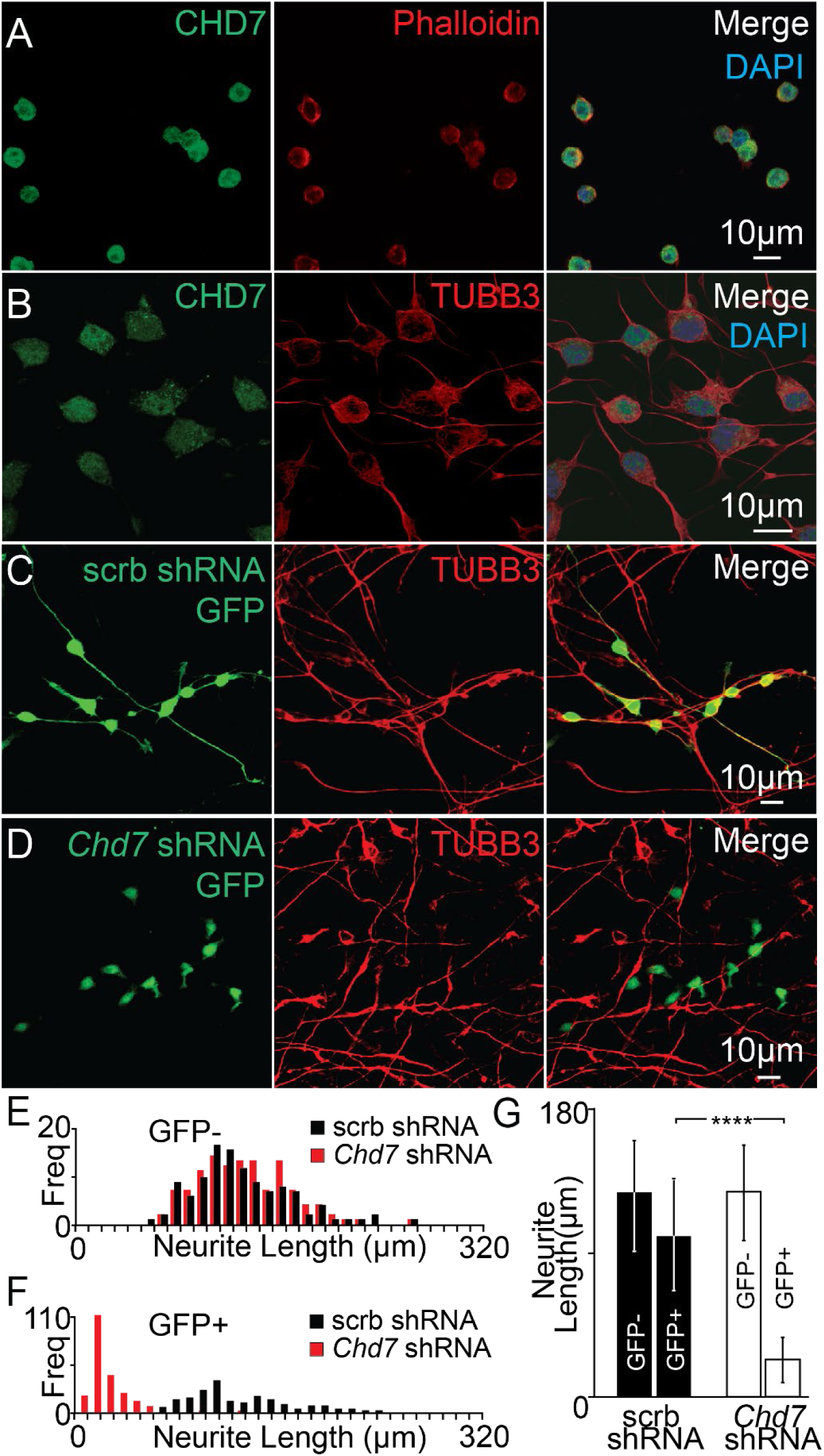
Knockdown of *Chd7* in iMOP cells undergoing neuronal differentiation. CHD7 expression in (A) phalloidin-marked progenitors and (B) TUBB3-labeled neurons. (C) Neurons were marked by GFP and TUBB3 after transfection with a plasmid transcribing a scrambled (scrb) shRNA and expressing a GFP reporter or (D) after transfection with a plasmid containing *Chd7* shRNA and a GFP reporter. (E) Distribution of neurite lengths in untransfected (GFP-) cells from scrb shRNA (black) (n=200 cells) or *Chd7* shRNA (red) (n=200 cells) cultures were used as internal controls. (F) Distribution of the neurite lengths from transfected (GFP+) cells in scrb shRNA (black) (n=200 cells) or *Chd7* shRNA (red) (n=200 cells). (G) Quantification of neurite lengths of control untransfected (GFP-) cells and transfected (GFP+) cells in scrb shRNA (n=6) and *Chd7* shRNA independent cultures (n=6). The Student’s t-test was used for statistical analysis between GFP+ cells transfected with scrambled or *Chd7* shRNA, **** p< 0.0001.

To control for variations in culture conditions during neuronal differentiation, TUBB3 labeled neurites of GFP-cells from scrambled shRNA and *Chd7* shRNA transfected cultures were measured and used as a criterion to ensure that culture conditions for neuronal differentiation were similar between samples. As internal controls, GFP-TUBB3+ cells from scrambled shRNA and Chd7 shRNA transfected cultures were used. A similar distribution of neurite lengths was observed, suggesting that neuronal differentiation conditions were comparable between cultures (Fig. 2E). To quantify the effects of *Chd7* knockdown, the lengths of neurites marked by GFP-expressing cells were measured in scrambled shRNA and *Chd7* shRNA transfected cultures. *Chd7* shRNA-transfected cells showed a dramatic reduction in neurite length compared to scrambled shRNA-transfected cells (Fig. 2F). Average neurite lengths for GFP-cells after transfection with scrambled shRNA or *Chd7* shRNA were 125.7+/-37.1 and 128.6+/-35.3 µm, respectively. Average neurite lengths of GFP+ cells after transfection with scrambled shRNA or *Chd7* shRNA were 101.1+/-37.1 and 23.6+/-16.4 µm, respectively. Comparison of GFP+ cells from scrambled and *Chd7* shRNA transfected cultures showed a significant 4.3-fold decrease in neurite lengths (p<1x10^-4^) after *Chd7* knockdown (Fig. 2G). These data suggest that culture conditions are consistent and comparable, showing that CHD7 is required for neuronal differentiation in iMOPs.

### Identifying genome-wide CHD7-bound *cis*-regulatory elements

Next, to identify the molecular basis of CHD7 function, we sought to identify CHD7 binding sites in these cells. CHD7 is enriched at different *cis*-regulatory elements in various cell types (Sanosaka, Okuno et al. 2022). To determine the genome-wide binding of CHD7, CUT&Tag (Cleavage Under Targets and Tagmentation) was performed on progenitors and neurons, followed by deep sequencing. Sequence reads were mapped to genomic regions, and enriched regions were identified as binding sites for the protein of interest. To characterize CHD7 binding to different regulatory elements genome-wide, we used H3K4me3, EP300, and CTCF to mark promoters, enhancers, and insulators. Antibodies used for CUT&Tag are provided in Table S2. SEACR (Sparse Enrichment Analysis for CUT&RUN) was used for peak calling to identify statistically significant enrichment in different genomic regions (Meers, Tenenbaum et al. 2019) from H3K4me3, EP300, and CTCF CUT&Tag results. CHD7 signal +/- 5kb around the summit regions of H3K4me3, EP300, and CTCF peaks were displayed as heatmaps with associated profile plots (Fig. 3A). The quantified signal centered around +/- 5kb of the peak summits with box and whisker plots (Fig. 3B), showed a statistically significant increase in CHD7 enrichment in neurons compared to progenitors at all marked *cis*-regulatory sites. The previously reported cross-regulation of CHD7 on the *Sox2* gene aligns with the CUT&Tag results (Gao, Skidmore et al. 2024). Indeed, increased CHD7 enrichment in neurons was observed prominently at H3K4me3 (promoter) and EP300 (enhancer) sites around the *Sox2* gene (Fig. 3C). Motif analysis further confirmed SOX2 motifs with CHD7 CUT&Tag peaks in both progenitors (*p* = 1e^-67^) and neurons (*p* = 1e^-169^) (Fig. 3D). Since CHD7 associates with enhancers and promoters in a cell type-specific manner (Schnetz, Bartels et al. 2009), we identified genome-wide promoter and enhancer sites during otic neuronal differentiation.

**Fig. 3.**
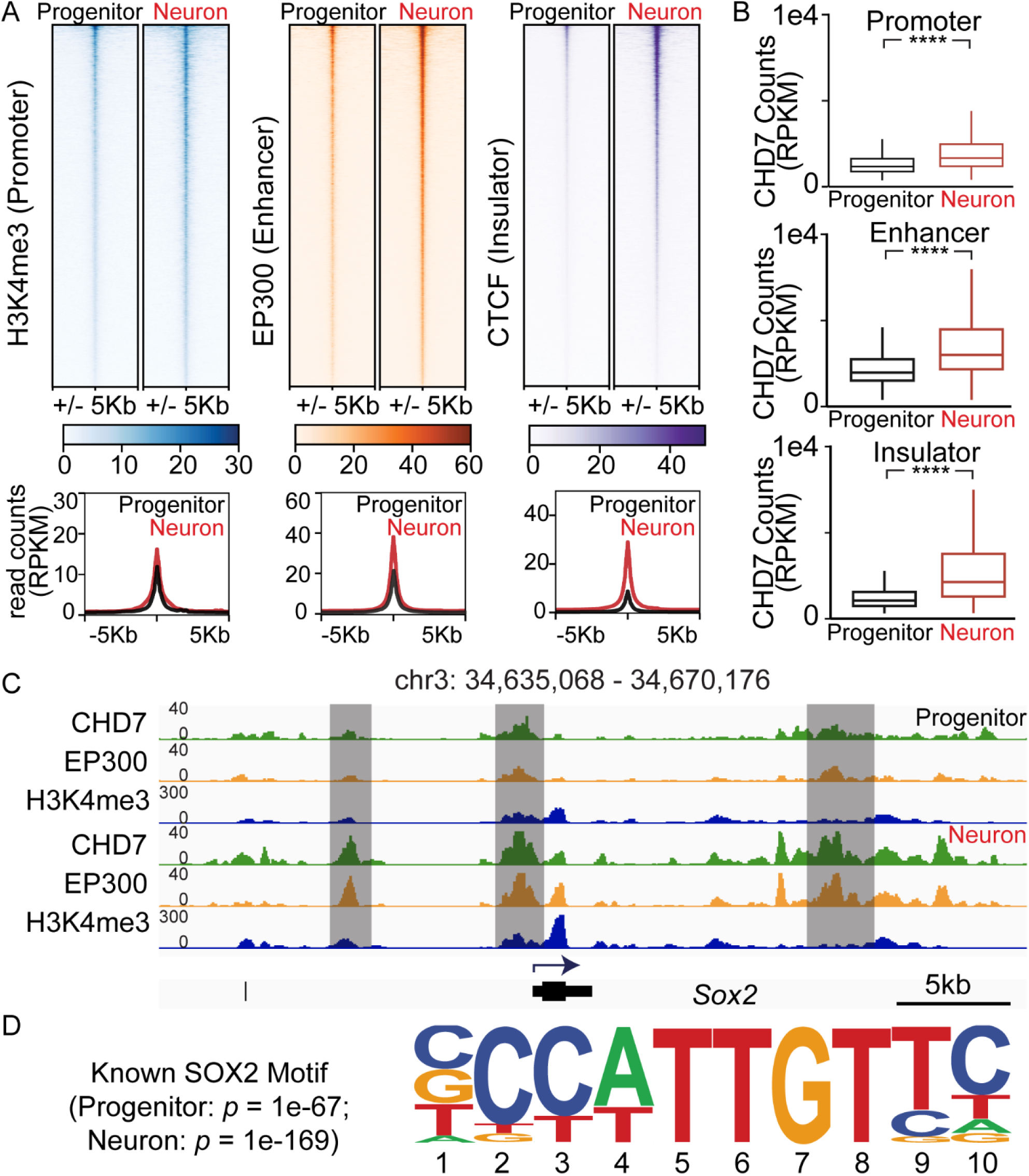
Correlating CHD7 sites to *cis*-regulatory marks. (A) Heatmaps and profile plots showed CHD7 reads within +/- 5kb windows of H3K4me3 (Promoter), EP300 (Enhancer), and CTCF (Insulator) in progenitors and neurons. (B) Box and whisker plots display normalized read counts (Reads Per Kilobase per Million, RPKM) for CHD7 at promoter, enhancer, or insulator (± 5kb) in neurons significantly increased compared to progenitors. (C) IGV (Integrative Genomics Viewer) displayed CHD7, EP300, and H3K4me3 enrichment at the *Sox2* gene in progenitors and neurons. Highlighted regions showed increased CHD7 signals in neurons. (D) HOMER motif analysis showed known SOX2 motifs in CHD7 SEACR peaks in both progenitors and neurons.

### CHD7 co-occupies CTCF binding sites

In sharp contrast to CHD7’s association with promoters and enhancers, little is known about CHD7’s function at insulators. During the analysis, we noticed that CHD7 enrichment at CTCF sites. CHD7 showed significant enrichment in neurons relative to progenitors, suggesting that CHD7 may function at different CTCF sites during neuronal differentiation. CTCF is a zinc finger protein that regulates gene expression by acting as a transcriptional activator or a transcriptional repressor, and it is also a well-known insulator-associated protein. CTCF is essential for cell fate transition, and deletion of CTCF prevents embryonic stem cells (ESCs) from differentiating into neural precursor cells (NPCs) (Kubo, Ishii et al. 2021). CTCF marks boundaries for topologically associating domains (TADs), and disruption of CTCF sites results in aberrant gene expression (Chang, Ghosh et al. 2023, Kim, Rahme et al. 2024). We hypothesize that regions marked by CHD7 and CTCF may function as insulators to regulate neuronal differentiation. To identify CHD7-bound insulators that facilitate the differentiation of progenitors into neurons, putative CHD7-marked insulators were first defined as CTCF and CHD7-bound regions (CTCF+CHD7+) in progenitors and neurons. Motif analysis using the CTCF+CHD7+ regions confirmed a statistically significant enrichment of the CTCF-binding motif in both progenitors (*p* = 1e^-176^) and neurons (*p* = 1e^-540^) (Fig. 4A). These data indicate that CHD7 is enriched at CTCF sites during neuronal differentiation.

**Fig. 4.**
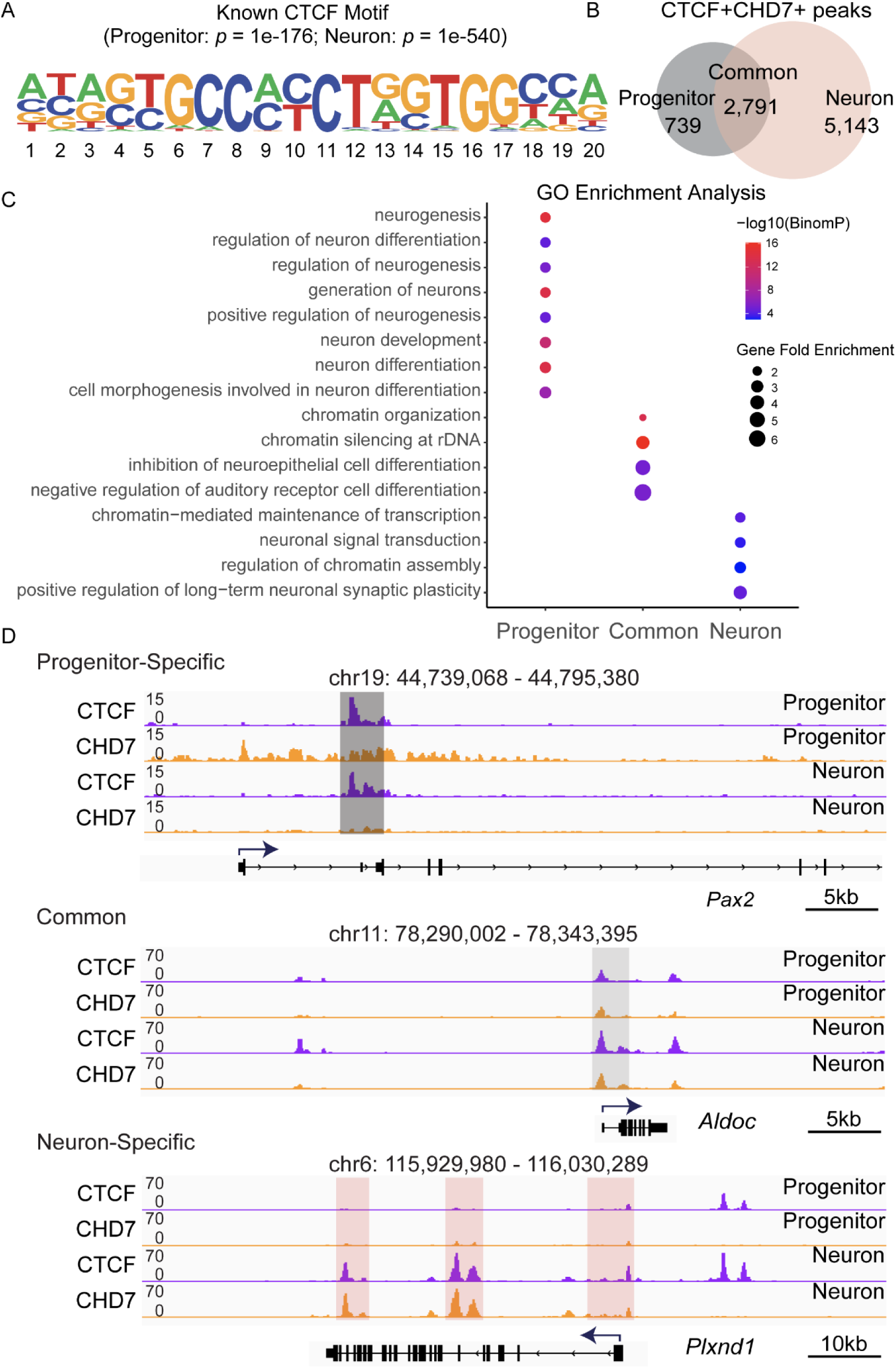
Identifying CTCF+CHD7+ regions near otic genes. (A) CHD7 bound insulators (CTCF+CHD7+) were defined as genomic regions marked by CTCF and CHD7. Motif analysis using HOMER showed known CTCF motifs in CTCF+CHD7+ SEACR peaks in both progenitors and neurons. (B) Venn diagram displayed the numbers of progenitor-specific, common, and neuron-specific CHD7 bound insulators. (C) The dot plot showed enriched GO (Gene Ontology) terms in progenitor-specific, common, and neuron-specific CHD7-bound insulators. (D) Enrichment of CTCF and CHD7 at progenitor-specific (*Pax2*), common (*Aldoc*), and neuron-specific (*Plxnd1*) genes. These genes were previously implicated in inner ear development and neuronal differentiation. Highlighted regions denote progenitor-specific (dark grey), common (grey), and neuron-specific (pink) CTCF+CHD7+ binding sites.

### Identifying cell type-specific CTCF+CHD7+ sites in progenitors and neurons

CTCF binding is known to be different in distinct cell types. To identify putative cell type-specific insulators marked by CHD7, we further categorized CTCF+CHD7+ peaks into three groups: regions that are only in progenitors (739 elements), only in neurons (5,143 elements), and shared regions (2,791 elements) (Fig. 4B). To understand the function of putative CHD7-bound insulators, CTCF+CHD7+ regions were used to perform functional enrichment using rGREAT (Gu and Hubschmann 2023). The functional enrichment analysis provided information about the biological processes associated with annotated CTCF+CHD7+ regions. The simplifyEnrichment package was used to visualize the terms related to the functional enrichment results and to reduce the complexity of the identified biological processes (Gu and Hubschmann 2023) (Fig. S3). A dot plot was used to visualize the biological terms associated with cell type-specific regions, where the size of the dot indicates the number of genes observed in the gene set from that biological process, while the color indicates the statistically significant p-value (Fig. 4C). Progenitor-specific CTCF+CHD7+ sites correlated to biological processes such as neuron differentiation, regulation of neuron differentiation and cell morphogenesis involved in neuron differentiation. Common CTCF+CHD7+ sites displayed terms such as inhibition of neuroepithelial cell differentiation and negative regulation of auditory receptor cell differentiation. In contrast, neuron-specific CTCF+CHD7+ regions displayed gene ontology enrichment terms such as neuronal signal transduction, regulation of chromatin assembly, and positive regulation of long-term neuronal synaptic plasticity (Fig. 4C). We interpret the results that progenitor CTCF+CHD7+ regions are involved in initiating neuronal differentiation. Meanwhile, neurons have distinct sites that maintain neuron-specific signaling pathways. The shared biological processes suggest that sensory epithelium and hair cell differentiation are actively inhibited as neuronal differentiation occurs. The results indicate that many identified CTCF+CHD7+ sites could regulate biological processes that facilitate neuronal differentiation.

Functional analysis suggested that cell type-specific CTCF+CHD7+ sites likely affect different gene regulatory networks at various stages of neuronal differentiation. We observed several genes involved in otic development and neuronal differentiation from the target genes. *Pax2* is an example gene near progenitor-specific CTCF+CHD7+ sites. *Pax2* is essential for otic development, and CHD7 has been implicated in regulating *Pax2* expression during early otic development (Gao, Skidmore et al. 2024). We observed both CTCF and CHD7 enrichment at *Pax2* in progenitors (Fig. 4D). Of the common CTCF+CHD7+ sites, we observed peaks at *Aldoc* (Fig. 4D). *Aldoc* was reported to be expressed in the inner ear (Fujita, Aoki et al. 2014, Sanders and Kelley 2022). Of the neuron-specific sites, we observed differential enrichment at *Plxnd1* (Fig. 4D)*. Plxnd1* was highly enriched in developing spiral ganglion neurons (Lu, Appler et al. 2011). The data suggests that CTCF+CHD7+ sites may regulate known otic genes during neuronal differentiation.

### Identifying CTCF+CHD7+ sites at TAD boundaries

While we observed numerous CTCF+CHD7+ sites near genes involved in otic and neuronal differentiation, we sought to determine how these co-bound regulatory regions might function as insulators to influence gene expression. Insulator elements, marked by CTCF, are often located at the boundaries of TADs, the fundamental units of 3D genome organization. CTCF plays a crucial role in defining TAD boundaries, which are largely conserved across cell types and species (Dixon, Selvaraj et al. 2012, Rao, Huntley et al. 2014). Therefore, we utilized published Hi-C data from neural progenitor cells (NPC) to define TAD boundaries in our progenitor cells (Bonev, Mendelson Cohen et al. 2017). The TADs allowed us to investigate the genomic context of the CTCF+CHD7+ sites. We defined TAD boundaries as a 40-kb region spanning ±20 kb from the reported TAD boundary coordinates from NPCs. Our analysis revealed multiple boundaries with CTCF+ CHD7+ sites. We focused on clusters of sites within the TAD boundary since they likely correspond to insulators. We focused on these regulatory elements, such as the ones marked as CTCF+CHD7+ sites 1-5 (Fig. 5A). Our findings suggest the presence of CHD7 at CTCF-marked insulators located within TAD boundaries.

**Fig. 5.**
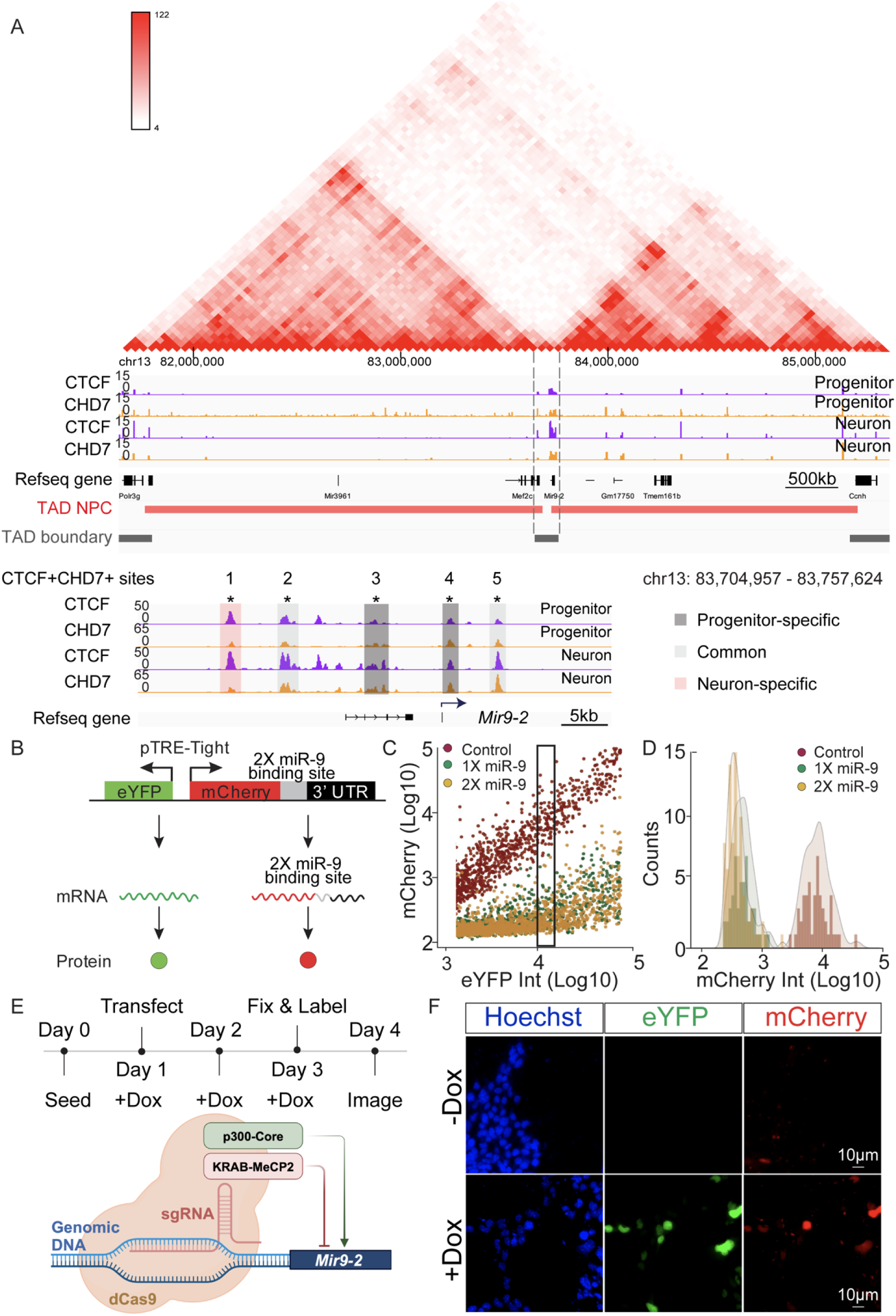
Identifying CTCF+CHD7+ sites at a TAD boundary and establishing a *Mir9-2* reporter. (A) CUT&Tag tracks display the enrichment of CTCF (purple) and CHD7 (orange) at specified genomic regions around the *Mir9-2* gene in progenitors and neurons. Published Hi-C data from neural progenitor cells (NPCs) were used to visualize topologically associating domains (TADs) (red triangle), TAD genomic regions (red bars), and TAD boundaries (black bars). Dashed lines highlight the TAD boundary around the *Mir9-2* gene. The bottom panel shows a zoom-in of the TAD boundary region annotated with CTCF and CHD7 signals. Highlighted signal tracks denote progenitor-specific (dark grey), common (grey), and neuron-specific (pink) CTCF+CHD7+ enriched genomic locations. sgRNAs were designed to target the CTCF+CHD7+ marked sites. (B) A dual fluorescent reporter expressing eYFP and mCherry was used to measure changes in *Mir9-2* gene expression, with mCherry containing two copies of miR-9 binding sites. (C) To determine the effects of miR-9 binding, 1 or 2 miR-9 binding sites were inserted between mCherry and the 3’UTR region. In the control, the 3’UTR after the mCherry gene was unmodified. Fluorescence of mCherry and eYFP from the reporter plasmid containing 0 (control) (red) (n=4), 1 (1X) (green) (n=4), and 2 (2X) (yellow) (n=4) copies of miR-9 binding sites in neurons. From the combined experiments, a total of 530 control, 619 1X miR-9, and 957 2X miR-9 cells were acquired. The eYFP fluorescence intensity was used to bin cells with similar transcriptional activity. Rectangle depicted an example of a bin used for analysis. (D) Histogram and associated density plots from a single bin showing the distribution of mCherry fluorescence in cells transfected with control, 1X, and 2X miR-9 reporter constructs. (E) The timeline outlined key experiments to investigate the effects of each CTCF+CHD7+ regulatory element on *Mir9-2* levels. Doxycycline (Dox) was added to the culture medium to express fluorescent proteins. Schematic depicts the CRISPRi (interference) and CRISPRa (activation) system. (F) Representative images showing the expression of eYFP and mCherry in transfected CRISPRi cells with or without Dox. Nuclei were labeled with Hoechst.

We surveyed genes within the TAD boundary to test whether perturbing the CTCF+CHD7+ sites affects gene expression. Genes that reside at these boundaries occupy a unique position at the interface between distinct regulatory domains, making them particularly sensitive to disruptions of the TAD structure (Lupianez, Kraft et al. 2015). While complete disruption of TAD structure could affect genes throughout the region, localized perturbations of TAD boundaries can significantly impact genes positioned directly at the boundary (Nora, Goloborodko et al. 2017, Chen, Ren et al. 2024, Kabirova, Ryzhkova et al. 2024). Disruption of a TAD boundary can compromise insulator function, lead to ectopic activation by enhancers in neighboring TADs, or a loss of local enhancer activation of genes (Tarjan, Flavahan et al. 2019, Xiao, Hafner et al. 2021). Altered TADs can significantly change gene expression at the boundary. In contrast, genes located deep within TADs are shielded from these effects and are primarily regulated by elements within their own TAD. They are less likely to experience dramatic changes in expression following localized boundary disruption.

Our functional analysis of CTCF+CHD7+ regions identified genes within the boundary and in the proximity of the putative insulators. One of the genes, *Mir9-2*, satisfied the criteria. The regulatory elements marked by CTCF and CHD7 around *Mir9-2* are evolutionarily conserved (Yuva-Aydemir, Simkin et al. 2011). *Mir9-2* has a well-documented role in neurogenesis, and the miR-9 family of microRNAs have been implicated in different aspects of neurogenesis (Coolen, Katz et al. 2013). miR-9 plays a key role in regulating the timing of neurogenesis by establishing a progenitor cell state that can either be maintained through self-renewal or progress toward neuronal differentiation (Coolen, Thieffry et al. 2012). We hypothesize that putative CHD7-marked insulators regulate *Mir9-2* levels to affect the transition from progenitors into neurons. We observed a cluster of CTCF+CHD7+ peaks near the *Mir9-2* gene (Fig. 5A). The CTCF+CHD7+ sites displayed different cell state enrichment and harbored progenitor-specific, shared, and neuron-specific sites. This finding implicates CTCF+CHD7+ marked regions at the TAD boundary may be essential for neuronal differentiation.

### Effects of perturbing CTCF+CHD7+ sites on *Mir9-2* levels

To determine the effects of the identified CTCF+CHD7+ sites in the TAD boundary, we used *Mir9-2* levels to indicate altered transcription. A bidirectional promoter driving transcription of enhanced yellow fluorescent protein (eYFP) and monomeric Cherry fluorescent protein (mCherry) was used (Mukherji, Ebert et al. 2011, Schmiedel, Klemm et al. 2015). Either one or two miR-9 binding sites were inserted at the 3′ untranslated region (3′UTR) of mCherry (Fig. 5B). Due to the known variability in transcriptional activity in the cells (Eling, Morgan et al. 2019), the fluorescence intensity of eYFP was used to define levels of transcriptional activity and to divide cells into bins for analysis. The mCherry transcript containing two miR-9 binding sites allows miR-9*-*dependent transcript degradation to affect mCherry protein expression. The mCherry fluorescence of cells from each bin was used as a read-out for miR-9 transcript levels (Fig 5B). To ensure that the dual fluorescence reporter containing miR-9 binding sites reflects miR-9-mediated transcript degradation, constructs containing no miR-9 binding site (control) or constructs containing 1 (1X miR-9) or 2 (2X miR-9) copies of miR-9 binding sites were transfected into cells. The eYFP and mCherry fluorescence intensities were acquired, and fluorescence intensities from individual cells were plotted. In neurons when *Mir9*-2 was upregulated, cells transfected with 1X or 2X miR-9 constructs displayed a significant reduction of mCherry signal compared to control (Fig. 5C). To ensure cells have comparable transcriptional profiles, eYFP fluorescence intensities were divided into bins to compare cells with similar transcriptional activities. The mCherry fluorescence signal from cells was obtained and plotted within each bin as a histogram. The mean fluorescence intensity of mCherry and the standard deviations for reporter constructs were calculated for each bin. The presence of miR-9 in the neurons significantly reduced the mCherry transcript and protein fluorescence intensities in bins where the eYFP levels were comparable (Fig. 5D). The 2x miR-9 reporter displayed less variance and was used to monitor the miR-9 levels.

To determine the function of individual CTCF+CHD7+ regulatory element functions, we generated stable CRISPRi (interference) iMOP cells expressing dCas9-KRAB-MeCP2 and stable CRISPRa (activation) iMOP cells expressing dCas9-p300-core. The CRISPRi and CRISPRa iMOP cells were transfected with a plasmid harboring the single guide RNA (sgRNA) and doxycycline-inducible 2X miR-9 reporter along with a plasmid expressing the reverse tetracycline-controlled transactivator (rtTA). Doxycycline binds rtTA and induces eYFP and mCherry expression. Cells were fixed and harvested for analysis. The CRISPRi and CRISPRa systems allowed the manipulation of the *cis*-regulatory elements around *Mir9-2* while simultaneously assaying for miR-9 transcript levels (Fig. 5E). In the absence of doxycycline, eYFP or mCherry was not detected in Hoechst-labeled cells. In contrast, adding doxycycline resulted in robust expression of both fluorescent proteins (Fig. 5F).

Using the CRISPRi and CRISPRa systems, individual plasmids that harbored sgRNAs were transfected into cells to target each CTCF+CHD7+ site. An empty plasmid that did not contain the sgRNA was used as a control. The eYFP and mCherry fluorescence were acquired, and the fluorescence intensities from both fluorescent proteins were measured from each Hoechst-labeled cell. eYFP-expressing cells were placed into 5 bins based on their fluorescence intensities. The mCherry fluorescence was an inversely correlated measure of *Mir9-2* levels. Using CRISPRi, we interfered with the CTCF+CHD7+ regions defined as progenitor-specific, common, and neuron-specific using sgRNA. We did not observe changes in mCherry fluorescence in any bins when using sgRNA1 (Fig. 6A). Similarly, no significant changes were observed for sgRNA2 and sgRNA3 (Fig. S4A, B). These results suggest that the regions are not active insulators for neuronal differentiation. We used these regions as negative controls. However, we observed a significant decrease in mCherry fluorescence across all eYFP bins after introducing sgRNA4 (Fig. 6B) and sgRNA5 (Fig. 6C). The data suggests that interfering with the two of the CTCF+CHD7+ sites decreased mCherry fluorescence by increasing *Mir9-2* transcription.

**Fig. 6.**
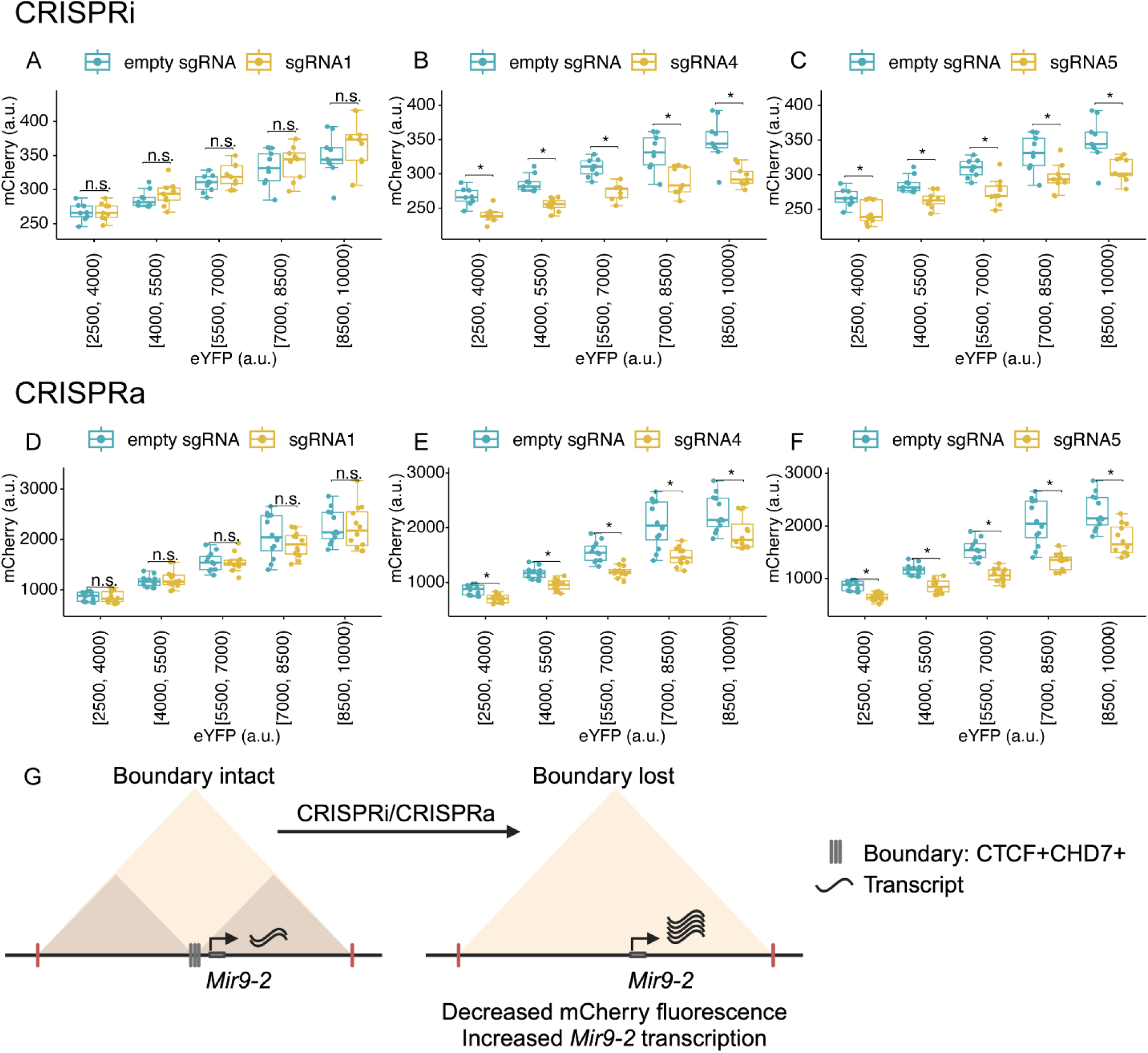
Blocking CTCF+CHD7+ around the *Mir9-2* gene during neuronal differentiation. (A-C) Fluorescence of mCherry and eYFP from inducible dual fluorescent reporter plasmid containing no sgRNA (empty sgRNA) and individual sgRNAs in dCas9-KRAB-MeCP2 iMOP cells undergoing early-stage neuronal differentiation (n=3, independent experiments). Box plots were used to compare mCherry levels between control and individual sgRNAs across different eYFP fluorescence bins. sgRNA1 didn’t show significant changes in mCherry level compared to empty sgRNA. sgRNA4 and sgRNA5 showed a significant decrease in mCherry level compared to the control empty sgRNA across different eYFP fluorescence bins. (D-F) Fluorescence of mCherry and eYFP from inducible dual fluorescent reporter plasmid containing no sgRNA (empty sgRNA) and individual sgRNAs in dCas9-p300-core cells undergoing early-stage neuronal differentiation (n=4, independent experiments). sgRNA1 didn’t show significant changes in mCherry level compared to empty sgRNA. sgRNA4 and sgRNA5 showed a significant decrease in mCherry level compared to the control empty sgRNA across different eYFP fluorescence bins. (G) Proposed model showing how CHD7 may function at an insulator to regulate gene expression. *The Mir9-2* gene is located within the TAD boundary, where there are multiple CTCF+CHD7+ binding sites. Normally, CHD7 is enriched at the insulators to inhibit *Mir9-2* gene expression. Disruption of the CTCF+CHD7+ sites by dCas9 complexes (CRISPRi and CRISPRa) increases *Mir9-2* gene expression.

Next, we used CRISPRa to activate the CTCF+CHD7+ regions to perform the converse experiment. No significant changes in mCherry fluorescence were observed for sgRNA1 (Fig. 6D), and the region was used as a negative control. Targeting with sgRNA2 and sgRNA3 showed a significant decrease in mCherry fluorescence in some bins (Fig. S4C, D). The results suggest that these regions may serve as weak insulators. Using CRISPRa, we observed a significant decrease in mCherry fluorescence across all eYFP bins after introducing sgRNA4 (Fig. 6E) and sgRNA5 (Fig. 6F) compared to the empty sgRNA control. Given that both CRISPRi and CRISPRa decreased mCherry fluorescence at CTCF+CHD7+ regions targeted by sgRNA4 and sgRNA5, these regions serve as strong insulators in the boundary region. We propose that CHD7 is required for the insulators to maintain TAD architecture to prevent promiscuous promoter-enhancer interactions between TADs. We prevented insulators from properly functioning when we blocked protein binding at the CTCF+CHD7+ sites within the TAD boundary, regardless of the CRISPR method. The inhibition of insulator function increased promiscuous enhancer interaction with the *Mir9-2* promoter, thus increasing *Mir9-2* transcript levels (Fig. 6G).

## DISCUSSION

### Identification of CHD7-bound insulators

Chromatin remodeling proteins play a crucial role in shaping the regulatory landscape of the genome, influencing gene expression and ultimately dictating cell identity. We performed genome-wide profiling of CHD7, a nucleosome repositioning enzyme, in immortalized multipotent otic progenitor (iMOP) cells, revealing its association with different *cis*-regulatory elements during neuronal differentiation. We observed CHD7 enrichment at promoters, enhancers, and insulators in both progenitor and neuronal stages, with a notable increase in CHD7 binding at these elements as progenitors differentiate into neurons. The increased CHD7 occupancy suggests a critical role for this nucleosome repositioner in establishing and maintaining the appropriate epigenetic landscape during neuronal differentiation. CHD7 can bind to short linker DNA but does not have a consensus DNA binding domain. However, CHD7 may interact with transcription factors to recruit it to specific chromatin sites. Our immunoprecipitation studies showed that CHD7 biochemically interacts with SOX2, similar to observations *in vivo*. The results suggest that iMOPs may serve as a cellular system to study gene regulatory networks in spiral ganglion neuron development. In human neural crest cells, CHD7 was associated with a polybromo- and BRG1- associated factor-containing complex implicated in directing neural crest development (Bajpai, Chen et al. 2010, Li, Xiong et al. 2013). Co-occupancy of CHD7 and CTCF genome-wide suggests that CHD7 may be part of a CTCF protein complex or be recruited by another DNA binding protein to these sites, as illustrated by the enrichment of CTCF and CHD7 at genes involved in otic development and neuronal differentiation.

Changes in histone modification and nucleosome repositioning at promoters and enhancers are well established. Here, we identified the association of CHD7 with regulatory elements, such as promoters and enhancers that facilitate otic neuronal differentiation. In addition, we showed that CHD7 also co-occupied CTCF sites. CTCF enrichment is a hallmark of insulator elements (Kim, Abdullaev et al. 2007). Insulators act as barriers to prevent the spread of chromatin state and prevent inappropriate enhancer-promoter interactions. Insulators ensure enhancers activate only their intended promoters to regulate target gene expression. CTCF sites are instrumental in forming TADs, the fundamental units of 3D chromatin organization within the nucleus (Nanni, Ceri et al. 2020). The interaction between CTCF and insulators is crucial for chromatin loop formation, enhancer-promoter insulation, and maintaining TAD boundaries. This coordinated action ensures precise gene regulation during neuronal differentiation. CTCF and CHD7 enriched sites also correlated to biological processes that prevent hair cell differentiation and facilitate neurogenesis when subjected to gene ontology analysis. Many surveyed genes were implicated in inner ear development and neuronal differentiation. Together, our results implicate that CHD7 chromatin remodeling is essential for the function of CTCF-marked insulators that facilitate neuronal differentiation.

### Perturbing CTCF+CHD7+ sites

TADs compartmentalize the genome and facilitate promoter-enhancer interactions within a single TAD while preventing interactions between promoters and enhancers in separate TADs. Therefore, disruption of TAD boundaries can lead to aberrant gene expression due to inappropriate promoter-enhancer communication (Lupianez, Kraft et al. 2015). Based on the co-enrichment of CHD7 and CTCF, we identified candidate TAD boundary sites in iMOP cells. We showed that CRISPRa and CRISPRi perturbations at the TAD boundary sites lead to increased gene expression of *Mir9-2*, a conserved microRNA involved in neurogenesis. We interpret that occluding the insulator sites with dCas9 prevents the binding of proteins such as CHD7 and CTCF at these sites and renders the insulator dormant. We suggest that CHD7 establishes the chromatin state at insulators and modulates the functional activity of TAD boundaries to maintain nuclear architecture for proper gene regulation.

### Identification of CHD7-bound insulators between TADs

TAD boundaries are often characterized by clusters of divergent CTCF binding sites, which function as insulators. These divergent CTCF sites at insulators interact with CTCF sites located far away, often in convergent orientations, facilitating the formation of chromatin loops. This interaction brings distant DNA regions closer to establish and maintain TADs. The orientation and interaction of CTCF-bound sites are crucial for the genome’s spatial organization, ensuring proper gene regulation by insulating regulatory elements and maintaining distinct regulatory domains. The clustering of CTCF sites with convergent orientations and other factors like cohesin and Mediator creates a robust boundary that restricts the influence of enhancers and other regulatory elements to their appropriate TAD compartment. The presence of CHD7 at CTCF-bound sites suggests that it could be involved in modulating boundary regions by influencing CTCF binding, cohesin loading, or other aspects of chromatin organization at these critical loci. Perturbing the boundary element by either *Chd7* shRNA knockdown or, more specifically, blocking the CTCF+CHD7+ sites allows promiscuous interaction of enhancers with promoters between TADs to dysregulate gene expression. We showed that expression of the well-conserved microRNA, *Mir9-2,* increased when the CHD7-bound insulators were masked. Our findings implicate that CHD7-bound insulators are critical in establishing gene expression that facilitates neuronal differentiation.

### Effects of perturbing CHD7-bound insulators on *Mir9-2* levels

*Mir9-2* is a well-conserved microRNA involved in neurogenesis. After cleavage and processing of the precursor *miR9-2* transcript, the mature miR-9 destabilizes target mRNAs to decrease transcript levels. In addition to repressing the expression of target genes (Baek, Villen et al. 2008, Selbach, Schwanhausser et al. 2008), many well-conserved miRNAs reduce noisy gene expression (Ebert and Sharp 2012, Schmiedel, Klemm et al. 2015). miR-9, in particular, has been shown to affect multiple transcripts that antagonize neurogenesis (Coolen, Thieffry et al. 2012).

The expression of multiple miRs during inner ear development has previously been described (Weston, Pierce et al. 2006, Elkan-Miller, Ulitsky et al. 2011, Rudnicki and Avraham 2012). The presence of miR-9 in the cochlea has been shown by qPCR (Weston, Pierce et al. 2006). Knockout of Dicer, an essential component of the miRNA processing complex, results in profound developmental deficits in the inner ear that cause spiral ganglion degeneration and hair cell innervation deficits by E17.5 (Friedman, Dror et al. 2009, Soukup, Fritzsch et al. 2009). In the brain, members of the miR-9 family are abundantly expressed (Lagos-Quintana, Rauhut et al. 2002, Kloosterman, Wienholds et al. 2006). A murine knockout of *miR9*-2 and *miR9*-3 leads to reduced production of early-born neurons within the pallium of the developing telencephalon, indicating that miR-9 plays a crucial role in neuronal differentiation in the telencephalon (Shibata, Nakao et al. 2011). Although much attention has been given to miR targets, the regulation of miRs in the inner ear has essentially been left unexplored. Here, we show that CHD7 binds to insulators around the *Mir9-2* gene and maintains the appropriate transcription of *Mir9-2* during neuronal differentiation. Changes in miR-9 levels can drastically alter the mRNA repertoire. miR-9 inhibited progenitor-specific genes that antagonize differentiation (Cao, Pfaff et al. 2007, Coolen, Thieffry et al. 2012). Continued expression of progenitor genes due to reduced miR-9 levels would prevent differentiation. The idea is supported by a previous report showing that murine CHD7 was required to maintain the appropriate number of delaminating neuroblasts during inner ear neurogenesis (Hurd, Poucher et al. 2010). Without CHD7, dysregulation of *Mir9-2* levels may contribute to improper SGN development in the inner ear. We propose that the CHD7-*Mir9-2* axis contributes to the development of SGNs.

Our study provides mechanistic insights into how CHD7 functions during SGN development by establishing the epigenetic state of insulator elements to maintain TADs, regulate 3D chromatin organization, and ultimately modulate global gene expression. We propose that decreased CHD7 activity may alter the nuclear organization and indirectly affect gene expression, resulting in improper differentiation and generation of SGNs. Our study also implicates that decreased CHD7 activity may prevent proper development and reduce the number of SGNs, leading to sensorineural hearing loss observed in patients with CHARGE syndrome.

## EXPERIMENTAL PROCEDURES

### iMOP Cell Culture

iMOP cells were grown in suspension with DMEM/F12 (Life Technologies) containing 1XN21 supplement (Life Technologies), 25 µg/ml carbenicillin, and 20 ng/ml bFGF (PeproTech). Cells were dissociated and cultured for 3 days before dissociation and plating onto 10 µg/ml poly-D-lysine and 10 µg/ml laminin-coated surfaces for neuronal differentiation. Culture media was switched to neurobasal media (Life Technologies) containing 1XN21, 2 mM L-glutamine (Life Technologies), and Cdk2 inhibitor (Selleckchem). Media was changed every other day. Cells were maintained in neurobasal media for 1 to 7 days before being used for immunostaining or harvested for experiments. Cells were grown in 12 mm coverslips or a 96-well flat bottom plate for immunostaining. The coverslips were sterilized with 70% ethanol and exposed to UV for 20 minutes in the tissue culture hood. Dried coverslips were placed on wells in tissue culture plates and coated with 10 µg/mL of poly-D-lysine for 1 hour. The coverslips were washed 3 times in PBS and coated overnight with 10 µg/mL of laminin. The coverslips were washed 3 times with PBS, and cells were plated in the center of the coverslip and incubated at 37°C.

### RT-qPCR

Total RNA was extracted using Trizol reagent (Life Technologies) according to manufacturer instructions. According to manufacturer instructions, 1 µg RNA was used to make cDNA using the qScript cDNA synthesis kit (Quanta Biosciences). Relative levels of cDNA were measured by quantitative real-time PCR using SYBR green Taq polymerase (Life Technologies) for 40 cycles of 95°C for 15 s, 60°C for 1 min using the StepOnePlus real-time PCR machine. The relative transcript levels were normalized to *Gapdh* and compared to proliferating iMOP cells unless otherwise indicated. The ΔΔCT method was performed for all the normalization and comparisons. Primers used for qPCR are listed in Fig. S2D.

### Western Blot Analysis

Cells were lysed in lysis buffer (50 mM Tris/HCl, pH 7.5, 150 mM NaCl, 1 mM EDTA, 1 mM EGTA, 1% Triton X-100) containing 10% glycerol and containing phosphatase inhibitor (Thermo Scientific) and a mixture of protease inhibitors (Roche). Protein lysates (30 µg) were loaded and separated on 4-12% Bis-Tris Novax NuPAGE gradient gels (Life Technologies), transferred to PVDF membrane, and incubated in blocking buffer (phosphate-buffered saline (PBS), 0.1% Tween 20 and 5% nonfat dried milk) for 1 hour. Proteins of interest were detected by incubating membranes overnight at 4°C with primary antibodies. Immunoreactive bands were detected by incubating them with horseradish peroxidase-conjugated secondary antibodies and applying a chemiluminescence substrate (Pierce ECL, Thermo Fisher Scientific) or using fluorescently conjugated antibodies. Chemiluminescence was detected by exposing membranes to either X-ray film (RPI) or captured on the LiCore Western blot detection system. Quantification of the intensity from individual bands was done using ImageJ.

### Immunoprecipitation

iMOP cells were lysed as previously described (Jadali, Song et al. 2016) and sonicated 3 times for 15 seconds at 30% power. 250 µg of precleared protein lysates were incubated with 5 µg of SOX2 antibody (AB5603, Millipore) or rabbit IgG (Sigma) overnight, followed by incubation with protein G beads. Final protein lysates were loaded on 4-12% Bis-Tris Novax NuPAGE gradient gels, and a Western blot was performed.

### Immunofluorescence labeling

iMOP cells were fixed with 4% paraformaldehyde (PFA) in 1XPBS for 30 minutes at room temperature, followed by three washes with 1XPBS. Cells were permeabilized in wash buffer (1XPBS, 0.1% Triton-X100) for 10 minutes and then incubated in blocking buffer (1XPBS, 0.1% Triton-X100, 10% normal goat serum) for 1 hour at room temperature. Primary antibodies against CDKN1B, TUBB3, or CHD7 were diluted in a blocking buffer and incubated with iMOP cells overnight at 4°C. After incubation with primary antibodies, cells were washed three times with wash buffer and then incubated with various combinations of Hoechst (100 ng/ml), DAPI (1µg/ml), Alexa Fluor 568-conjugated Phalloidin, Alexa Fluor 488-, Alexa Fluor 568-, or Alexa Fluor 647-conjugated secondary antibodies in blocking buffer for 2 hours at room temperature in the dark. Cells were washed and mounted on slides with Prolong gold antifade mounting media (Life Technologies). Antibodies used for immunofluorescence labeling are listed in Table S2. Immunofluorescence images were obtained using a Zeiss LSM 800 confocal microscope with a 40X 1.3 NA water immersion objective or an Olympus DSU unit with a 60X 1.3 NA apochromatic oil immersion objective. To trace neurites of differentiating iMOP cells were immunostained with TUBB3 after transfection with constructs derived from pLKO.3G. Fluorescence images of GFP and TUBB3-labeled iMOP cells were acquired, and cells with TUBB3-labeled neurites that reside within the field of view were used for quantification. TUBB3-expressing cells were categorized as GFP+ and GFP-before using the ImageJ Neurite tracer plugin to determine neurite lengths for each population of cells.

### Stable and Transient *Chd7* Knockdown

To generate a stable *Chd7* knockdown cell line, single-stranded complementary oligos containing either scrambled or *Chd7* shRNA sequence (Fig. S2E) were 5’ phosphorylated with polynucleotide kinase and annealed together to form a double-stranded oligo containing restriction site overhangs for cloning into lentiviral vectors. For the lentiviral vector containing a blasticidin resistance cassette, pLKO.1-blast (Addgene #26655), double-stranded oligos were ligated into the AgeI and EcoRI restriction sites. For the lentiviral vector expressing GFP, pLKO.3G (Addgene #14748), double-stranded oligo was cloned into the EcoRI and PacI restriction sites. Viral constructs and packaging plasmids were transfected into 293FT cells using the calcium phosphate co-precipitation method. The supernatant was collected 48- and 64 hours post-transfection and combined. The virus was concentrated by PEG 6000 precipitation (Kutner, Zhang et al. 2009). For stable *Chd7* knockdown, iMOP cells were infected at a multiplicity of infection (MOI) of 5. Twenty-four hours after infection, cells were selected with 10 µg/ml blasticidin. Cells were harvested 72 hours after selection. For transient shRNA knockdown experiments, iMOP cells were plated for neuronal differentiation and allowed to recover for up to 24 hours before transfection with pLKO.3G *Chd7* shRNA or pLKO.3G scrambled shRNA using jetPRIME (Polyplus). Cells were fixed 72 hours after transfection and immunostained. A simple neurite tracer from ImageJ was used to trace and measure neurite lengths of TUBB3-positive cells.

### Bulk RNA-seq

Total RNA was extracted from 1-5x10^6^ proliferating or neuronal iMOP cells using 1ml of TRIzol Reagent (Thermo Fisher Scientific #15596026) following the manufacturer’s instructions. Total RNA was treated with DNaseI (NEB #M0303S), and then DNaseI was removed by phenol-chloroform extraction. Following the manufacturer’s instructions, the ribosomal RNA was depleted from 500 ng of DNase-treated total RNA using the NEBNext rRNA depletion Kit (NEB #E6310S). Following the manufacturer’s instructions, the RNA was used for stranded RNA-seq library preparation with the Takara SMARTer Stranded RNA-seq kit (Takara Bio #634838). The library size is 300 bp, including adaptors of average size. Indexed libraries were normalized by concentration and pooled together. Pooled libraries were sequenced using the Illumina Novaseq 6000 S4 platform (2x150bp). Sequence reads were mapped to reference mouse genome (mm10) using HISAT2 (Kim, Langmead et al. 2015). The SAM files were converted, sorted, and indexed to generate sorted BAM files using Samtools (Li, Handsaker et al. 2009). The featureCounts command from Rsubread was used to create the count matrix, and differential expression was calculated using DEseq2 (Love, Huber et al. 2014). The results were visualized using EnhancedVolcano (Blighe K 2024).

### CUT&Tag

According to the manufacturer’s instructions, the progenitors and neurons were used for CUT&Tag experiments using the CUT&Tag-IT Assay Kit (Active Motif, cat# 53160). 500,000 cells were used for each reaction. Cells were incubated with different primary antibodies targeting the protein of interest, while cells prepared with secondary antibodies only served as a control. Antibodies used for CUT&Tag are listed in Table S2. DNA fragments were amplified and cleaned using SPRI beads. Finally, DNA libraries were sequenced with the Illumina HiSeq 3000 platform with a 150 bp paired-end sequencing configuration.

CUT&Tag fastq files for EP300 and H3K4me3 were acquired from the Gene Expression Omnibus (GEO) database (GSE250033). The remaining CUT&Tag results were generated as described in (Kim, Martinez et al. 2024) and were sequenced with the Illumina HiSeq 3000 platform with 150 bp paired-end sequencing. Paired-end CUT&Tag reads were aligned to the reference genome (mm10) with Bowtie2 (Langmead and Salzberg 2012). Only mapped reads were retained with Picard (https://broadinstitute.github.io/picard/ 2019) and used for the downstream analysis. Samtools was employed to convert SAM output files into BAM files (Li, Handsaker et al. 2009). DeepTools was used to eliminate duplicated reads, and uniquely mapped reads were normalized as Reads Per Kilobase per Million (RPKM) in bigwig files with the bamCoverage command (Ramirez, Dundar et al. 2014). Bigwig files were converted into bedgraph files for peak calling using the UCSC bigWigToBedGraph command (Kent, Zweig et al. 2010). Peak calling was performed using Sparse Enrichment Analysis for CUT&RUN (SEACR) (Meers, Tenenbaum et al. 2019). Bedgraph files from CHD7, EP300, H3K4me3, and CTCF libraries were used as target files, while libraries prepared with only secondary antibodies served as a threshold for peak calling. BEDtools was employed to select peaks with at least 10% reciprocal overlaps between replicates to identify consensus peaks for downstream analysis (Quinlan and Hall 2010). CUT&Tag signal tracks around genes of interest were visualized using Integrative Genomics Viewer (IGV) (Robinson, Thorvaldsdottir et al. 2011). HOMER was used to perform motif analysis on CHD7 peaks in both progenitors and neurons (Heinz, Benner et al. 2010).

### Identification of CHD7-bound insulators

To identify the CHD7-bound insulator genome-wide, CTCF was used as an insulator marker, and CHD7-bound insulators were defined as CTCF+CHD7+ in progenitors and neurons. CTCF+CHD7+ were identified as regions where CTCF and CHD7 peaks displayed at least one base pair overlap. Bedtools intersect command was employed to identify cell type-specific CHD7-bound insulators by their presence in progenitors and absence in neurons, or vice versa. Common CHD7-bound insulators were identified based on their presence in progenitors and neurons. HOMER was used to perform motif analysis on CTCF+CHD7+ peaks in both progenitors and neurons (Heinz, Benner et al. 2010).

### Heatmaps and profile plots

Heatmaps and profile plots were generated using signal and region files acquired from CUT&Tag data. Signal files are average bigwig files from two replicates generated using bigwigCompare, while region files are the summit regions corresponding to the maximum signals of SEACR peaks. CHD7 signals at *cis*-regulatory elements were visualized using CHD7 bigwig as signal files, while the summit regions of H3K4me3 (promoter), EP300 (enhancer), and CTCF (insulator) SEACR peaks were used as region files. Heatmaps and profile plots were generated with DeepTools to visualize CHD7 signals centered around +/- 5kb of the summit regions of H3K4me3, EP300, and CTCF.

### Quantification of heatmap signals

DeepTools was employed to extract corresponding CUT&Tag reads displayed in the heatmaps to quantify heatmap signals at specified genomic regions. Total heatmap signals corresponding to normalized CUT&Tag reads (RPKM) within specified genomic regions were plotted in box and whisker plots to denote the medians and quartiles. The Wilcoxon rank-sum test was performed in R to determine if there are statistical differences between progenitors and neurons. Unless noted, the p values from the test were defined as ****p < 0.0001 to be significant.

### Gene ontology (GO) enrichment analysis

CTCF+CHD7+ progenitor-specific, common, and neuron-specific regions were analyzed using rGREAT and SimplifyEnrichment in Rstudio. Gene ontology terms with corresponding keywords were identified, and results were visualized using GOPlot or clusterProfiler to represent the p values and percent of genes detected in the gene ontology terms.

### Generation of stable CRISPRi and CRISPRa iMOP cell lines

Stable CRISPRi (dCas9-KRAB-MeCP2) cell line and CRISPRa (dCas9-p300-core) cell line were generated using lentiviral particles produced from either the Lenti_dCas9-KRAB-MeCP2 (Addgene #122205) or Lenti-EF1alpha-dCas9-p300_Blast (Addgene #192653). HEK293FT packaging cells were transfected with the Lenti plasmids, pRSV-Rev (Addgene #12253), pMDLg/pRRE (Addgene #12251), and pMD2.G (VSVG) (Addgene #12259) using jetOPTIMUS transfection reagent (Polyplus). The supernatant was collected 48- and 72 hours post-transfection. Proliferating iMOPs were infected with unconcentrated lentivirus. Forty-eight hours after infection, cells underwent stepwise increases up to 5 µg/ml blasticidin for selection. Cells were expanded and maintained in 2 µg/ml of blasticidin for 3 weeks to obtain stable CRISPRa and CRISPRi cell populations.

### Establishing the *Mir9-2* dual fluorescence reporter

A dual fluorescence reporter was used to measure *Mir9-2* transcript levels. iMOP-derived neurons were co-transfected with pTRETightBI-RY-0 (Addgene #31463) as control or pTRETightBI-RY-0 containing one or two copies of miR-9 binding site along with the FUW-M2rtTA plasmid (Addgene #20342) 5 days after initiating neuronal differentiation using jetPRIME transfection reagent (Polyplus). Transfection condition for iMOP-derived neurons was accomplished according to the manufacturer’s conditions. The transfection reagent from cultures was removed and replaced with fresh medium after 24 hours. Cells were cultured in a neural basal medium containing B27, 2 mM L-glutamine, and 1 µg/ml of doxycycline to induce the expression of fluorescent proteins. 48 hours after transfection, cells were fixed and used to acquire quantitative fluorescence imaging.

For fluorescence intensity quantification assay, iMOP-derived neurons were fixed with 4% formaldehyde for 20 minutes at room temperature and then washed 3 times for 10 minutes each using 1XPBS. Cells were then incubated in blocking buffer (5% goat serum, 0.1% Triton X-100 in 1XPBS) for 1 hour at room temperature and then incubated with primary antibody against TUBB3 at 4^◦^C overnight. After incubation with primary antibodies overnight, cells were washed 3 times for 15 minutes each using washing buffer and then incubated with secondary antibodies for 2 hours at room temperature. Hoechst and Alexa Fluor 647-conjugated secondary antibodies were diluted in the blocking buffer. After secondary antibody incubation, cells were washed 3 times, each for 15 minutes, using a washing buffer. Cells were then rinsed with 1XPBS, and mounted on slides with a Prolong Gold antifade mounting medium (Thermo Fisher). INCell Analyzer 6000 (GE Healthcare) was used to acquire quantitative fluorescence images. DAPI, FITC, dsRed, and Cy5 channels were used to acquire Hoechst, eYFP, mCherry, and TUBB3 signals. Images were acquired using a 40X 1.0 NA air objective and saved as 16-bit images. Quantitative fluorescence images were then analyzed using IN Carta software.

*Mir9-2* levels were monitored using a modified pTRETightBI-RY-0 (Addgene #31463). Plasmids containing miR-9 binding sites were generated by cutting pTRETightBI-RY-0 with ClaI and EcoRV and ligated to synthesized oligos with compatible overhangs that contain one copy and two copies of miR-9 binding site (TCATACAGCTAGATAACCAAAGA). Plasmids were transfected into iMOP cells cultured in µClear 96 well plates (Greiner) with 1 µg/ml of doxycycline-containing medium.

48 hours after transfection, cells were fixed with 4% formaldehyde and subjected to immunofluorescence labeling with TUBB3 antibodies and Hoechst. Quantitative fluorescence images from the cells were acquired using a 40X 1.0 NA air objecting using the InCell Analyzer 6000 (GE Healthcare), a high-content confocal microscope, and saved as 16-bit images. Background signal was established as fluorescence in untransfected cells labeled with secondary antibody. To quantify the fluorescent intensity of acquired images, Hoechst labeled nuclei were first identified and used to define a mask. Using a collar algorithm, the TUBB3 signal was used to define the cell body. In transfected cells, the nuclear mask was used to determine the fluorescence intensity of eYFP fluorescence and mCherry. Decision trees were generated to identify and categorize eYFP, mCherry, and TUBB3-expressing cells. The fluorescence intensity measurements were exported to R studio for analysis. In short, cells between 2,500 and 54,000 eYFP arbitrary fluorescence units (a.u.) were used for analysis. Individual cells’ eYFP and mCherry fluorescence intensities were displayed on a scatterplot.

### Targeting of CHD7-bound insulators by CRISPRi and CRISPRa

To inactivate each CTCF+CHD7+ binding site around *Mir9-2*, a CRISPRi stable iMOP cell line was used, and five single guide RNAs (sgRNAs) were designed for each site (Table S1). Each sgRNA was cloned with a U6 promoter into the pTRETightBI-RY-0 miR-9-2C plasmid using Gibson Assembly. Each plasmid was named based on the sgRNA. For example, pTRETightBI-RY-0 miR-9-2C_U6_sgRNA1 represents that the sgRNA1 was cloned into the pTRETightBI-RY-0 miR-9-2C plasmid. Proliferating iMOP cells were seeded for neuronal differentiation and allowed to recover for 24 hours before co-transfection. The pTRETightBI-RY-0 miR-9-2C plasmid harboring the sgRNA and FUW-M2rtTA plasmids were co-transfected into cells using jetPRIME transfection reagent. Transfection conditions for iMOP-derived neurons were performed according to the manufacturer’s conditions. Neuronal differentiation medium with transfection reagents was replaced with fresh neuronal differentiation media containing 1 µg/ml of doxycycline 8 hours after transfection. Cells were allowed to undergo neuronal differentiation for another 48 hours and then were fixed for immunostaining. The same paradigm and sgRNAs were used to activate CTCF+CHD7+ around *Mir9-2* using CRISPRa.

### Quantification of fluorescence intensity of miR-9 fluorescent reporter

After acquiring images, the INCarta software was used to process images to quantify the fluorescence intensity of the images. The Robust algorithm was used to identify and define Hoechst-labeled nuclei, and Cy5 was used to identify TUBB3-labeled soma as a whole cell mask. Each cell’s nuclei size and fluorescence intensities of each fluorescence channel were measured. A classifier was used to set the criteria for identifying viable and successfully transfected cells for analysis. The following criteria for viable cells were selected: medium nuclei size (25 ∼ 130 µm^2^), low Hoechst labeled nuclei arbitrary fluorescence units (a.u.) ≤ 9000 a.u. and eYFP ≥ 1000 a.u. After filtering the fluorescence intensity measurements through INCarta, the data were exported into Rstudio for analysis. Cells with an eYFP intensity between 2500 and 10000 a.u. were used to analyze mCherry expression levels. The eYFP intensity was used to divide cells into five bins (bin size = 1500 a.u.) to compare cells with similar transcriptional activity. To compare the mCherry expression of each experimental group with the control group that does not contain a sgRNA (empty sgRNA), the outliers from mean ± 2 standard deviation of the mCherry intensity in each bin were excluded in the analysis. An unpaired Student’s T-test was used to compare the mCherry intensity between the control (empty sgRNA) and each experimental group in each big bin. P values were defined as *p < 0.05.

### Statistical Analysis

All error bars in the data are expressed as +/- standard deviation (sd) of values obtained from independent experiments unless otherwise stated. The numbers (n) of independent experiments from separate cultures were listed for experiments. Technical triplicates were included in each experiment. Statistical tests were done in R version 4.4.3. Normal distribution of data was done using the Shapiro-Wilks test. The unpaired two-tailed Student’s t-test compared control and experimental means in normally distributed samples. If sample distribution was not normally distributed, the Wilcoxon rank sum non-parametric test was used to compare sample medians. The sample sizes for each experiment are noted in the figure legends. For all figures, p values are defined as: * p<0.05, ** p<1x10^-2^, *** p<1x10^-3,^ and **** p<1x10^-4^ unless otherwise stated.

## CONFLICT OF INTEREST

The authors declare that there is no conflict of interest.

## AUTHOR CONTRIBUTIONS

JQ, AJ, JN, EM, ZS, and KYK designed the experiments and interpreted and analyzed the data. JQ, JN, and KYK performed bioinformatics analysis. JQ and KYK wrote the manuscript.

## ACKNOWLEDGEMENTS

We thank Jenna Holland for her assistance in cloning the miR-9 reporter. KYK is supported by R01 DC018404. KYK acknowledges previous support from the Duncan and Nancy MacMillan Faculty Development Chair Endowment Fund, Busch Biomedical Research Grant, Rutgers Faculty Development Grant, CHARGE Foundation Grant, and R01 DC015000.

## Supplemental Figures

**Fig. S1.**
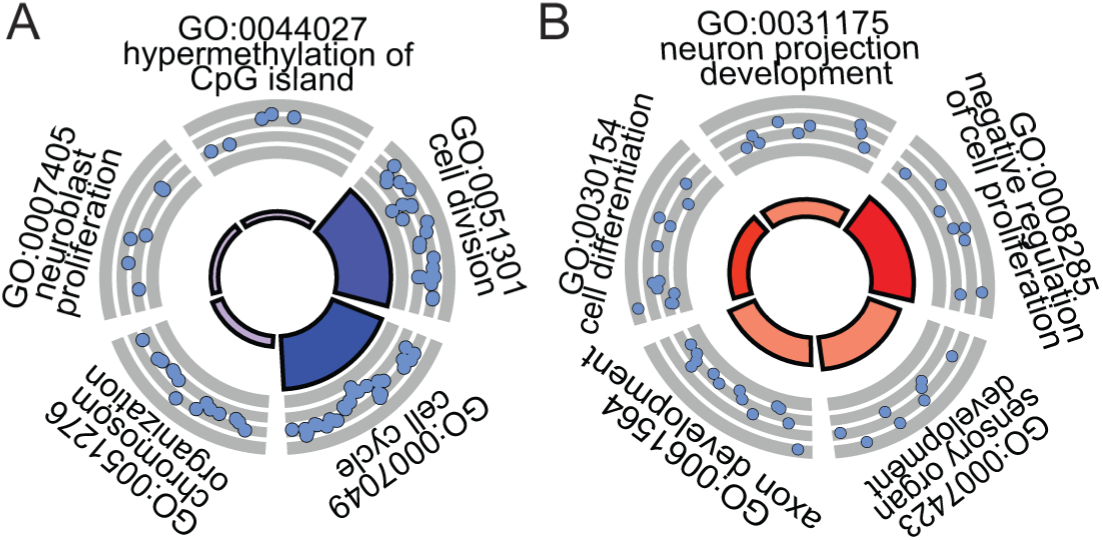
Gene Ontology (GO) analysis on differentially expressed genes in proliferating iMOPs and iMOP-derived neurons. Circos plot displaying select GO term descriptions with differential expression levels of genes within the GO process in (A) proliferating iMOPs and (B) iMOP-derived neurons.

**Fig. S2.**
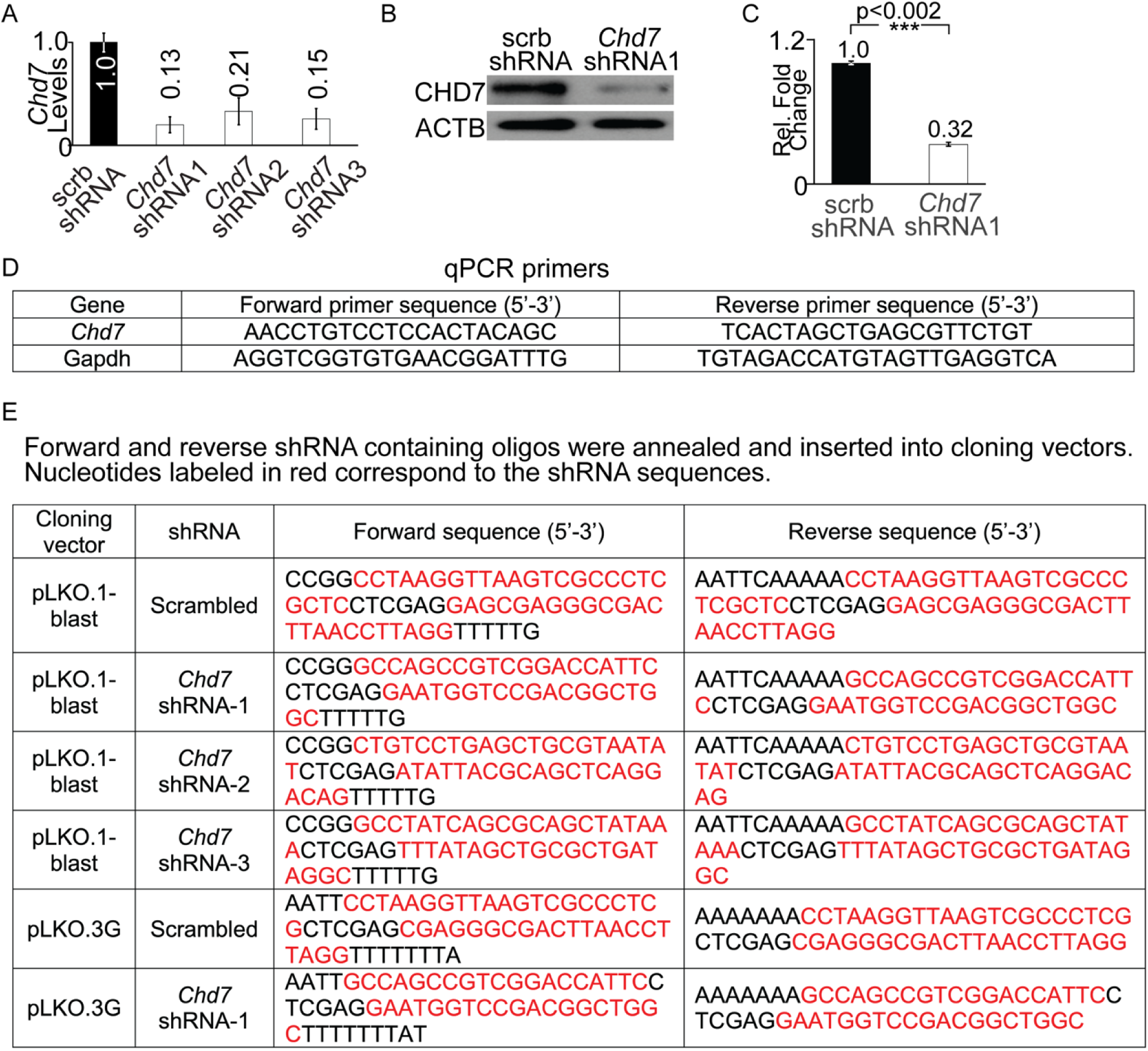
Knockdown of *Chd7* in iMOP cells. Differentiating iMOP cells were transduced with lentivirus containing scrambled (scrb) shRNA or three different *Chd7* shRNAs (*Chd7* shRNA 1-3). Cells were selected in a blasticidin-containing medium, and the total RNA was harvested from the cultures. (A) qPCR for *Chd7* transcript levels was performed from transduced cells and normalized to scrb shRNA control (n=3). (B) Representative Western blot of CHD7 expression levels in *Chd7* shRNA1 transduced cells using ACTB as a loading control (n=3). (C) Quantification of protein levels from Western blot showing a decrease in CHD7 to 0.32-fold (n=3) in *Chd7* shRNA1 compared to scrb shRNA. (D) qPCR primer sequences. (E) shRNA containing oligos that were annealed, extended and inserted into cloning vectors.

**Fig. S3.**
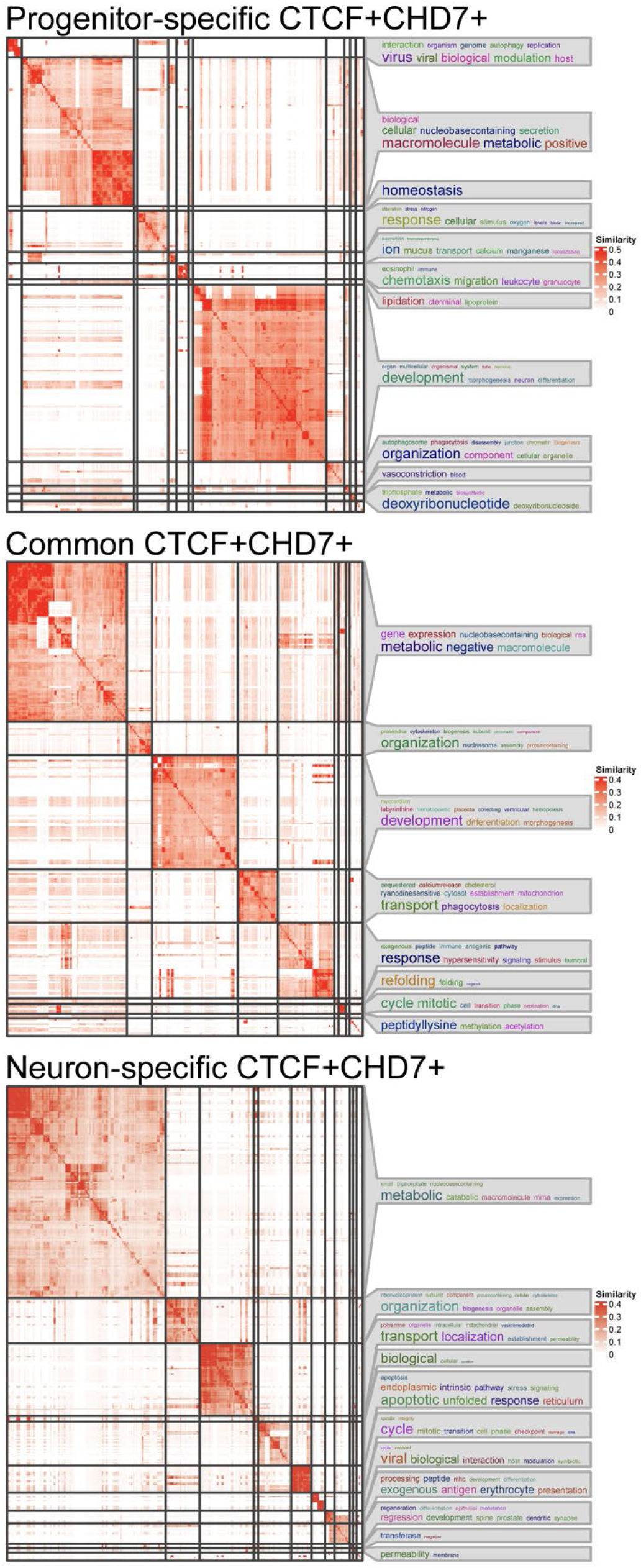
GO analysis on CTCF+CHD7+ enriched sites. GO terms associated with progenitor-specific, common, and neuron-specific CTCF+CHD7+ enriched sites. GO terms were clustered into similar groups and displayed as a matrix using the simplifyEnrichment package. Word clouds on the right of the matrixes display the key words that are over-represented in each category.

**Fig. S4.**
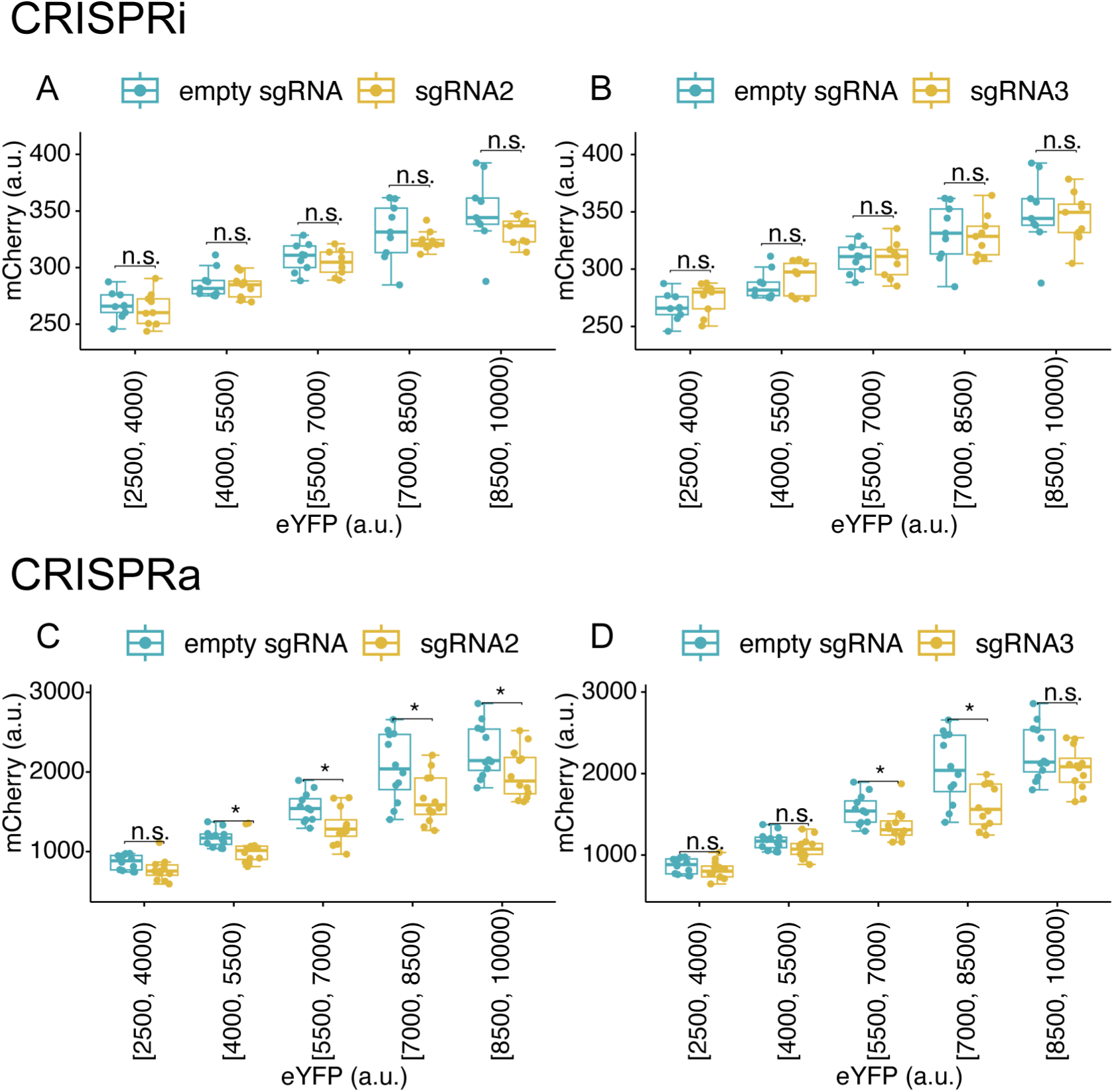
**Targeting CTCF+ CHD7+ regions around *Mir9-2*** (A-B) Fluorescence of mCherry and eYFP from inducible dual fluorescent reporter plasmid containing no sgRNA (empty sgRNA) and individual sgRNAs in dCas9-KRAB-MeCP2 iMOP cells undergoing early-stage neuronal differentiation (n=3, independent experiments). sgRNA2 and sgRNA3 did not show significant changes in mCherry level across different eYFP fluorescence bins compared to empty sgRNA. (C-D) Fluorescence of mCherry and eYFP from inducible dual fluorescent reporter plasmid containing no sgRNA (empty sgRNA) and individual sgRNAs in dCas9-p300-core cells undergoing early-stage neuronal differentiation (n=4, independent experiments). sgRNA2 and sgRNA3 showed a statistically significant decrease in mCherry level compared to the control empty sgRNA in some eYFP fluorescence bins.

**Table S1:**
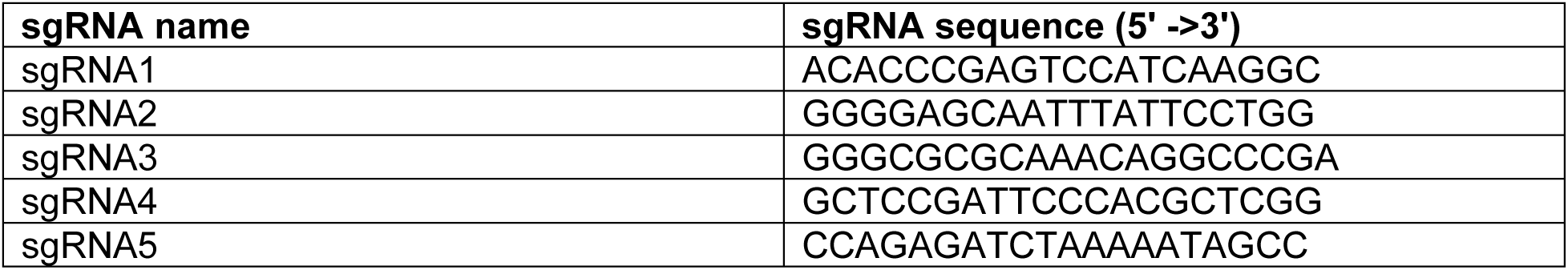
sgRNAs used for targeting regions around *Mir9-2* using CRISPRi and CRISPRa.

**Table S2.**

| **Reagent** | **Source** | **Identifier** | **Concentration /Dilution** | **Application** |
| --- | --- | --- | --- | --- |
| Mouse anti-TUBB3 | BioLegend | Cat# 801202, RRID:AB_2313773 | 1:1000 | Immunostaining |
| Rabbit anti-TUBB3 | BioLegend | Cat# 845501, RRID:AB_2566588 | 1:1000 | Immunostaining |
| Mouse anti-CDKN1B | Thermofisher/Lab Vision | Cat# MS-256-P RRID:AB_61871 | 1:500 | Immunostaining |
| Rabbit anti-CHD7 | Cell Signaling Technology | Cat# 6505, RRID:AB_11220431 | 1:300 | Immunostaining |
| Alexa Fluor 568 Phalloidin | Thermo Fisher Scientific | Cat# A12380 | 1:40 | Immunostaining |
| Goat anti-rabbit Alexa Fluor 488 | Thermo Fisher Scientific | Cat# A-11034, RRID:AB_2576217 | 1:3000 | Immunostaining |
| Goat anti-mouse Alexa Fluor 568 | Thermo Fisher Scientific | Cat# A-11031, RRID:AB_144696 | 1:3000 | Immunostaining |
| Goat anti-rabbit Alexa Fluor 647 | Thermo Fisher Scientific | Cat# A-21244, RRID:AB_2535812 | 1:3000 | Immunostaining |
| Rabbit anti-EP300 | Santa Cruz Biotechnology | Cat# sc-585, RRID:AB_2231120 | 1:50 | CUT&Tag |
| Rabbit anti-H3K4me3 | Active Motif | Cat# 39159, RRID:AB_2615077 | 1:50 | CUT&Tag |
| Rabbit anti-CHD7 | Cell Signaling Technology | Cat# 6505, RRID:AB_11220431 | 1:50 | CUT&Tag |
| Rabbit anti-CTCF | Active Motif | Cat# 61311, RRID:AB_2614975 | 1:50 | CUT&Tag |

## Notes

### Competing Interest Statement

The authors have declared no competing interest.

## REFERENCES

Ahmed, M., R. Moon, R. S. Prajapati, E. James, M. A. Basson and A. Streit (2021). "The chromatin remodelling factor Chd7 protects auditory neurons and sensory hair cells from stress-induced degeneration." Commun Biol 4(1): 1260.

Allen, M. D., T. L. Religa, S. M. Freund and M. Bycroft (2007). "Solution structure of the BRK domains from CHD7." J Mol Biol 371(5): 1135–1140.

Baek, D., J. Villen, C. Shin, F. D. Camargo, S. P. Gygi and D. P. Bartel (2008). "The impact of microRNAs on protein output." Nature 455(7209): 64–71.

Bajpai, R., D. A. Chen, A. Rada-Iglesias, J. Zhang, Y. Xiong, J. Helms, C. P. Chang, Y. Zhao, T. Swigut and J. Wysocka (2010). "CHD7 cooperates with PBAF to control multipotent neural crest formation." Nature 463(7283): 958–962.

Blighe K, R. S., Lewis M (2024). "EnhancedVolcano: Publication-ready volcano plots with enhanced colouring and labeling." https://github.com/kevinblighe/EnhancedVolcano.

Bonev, B., N. Mendelson Cohen, Q. Szabo, L. Fritsch, G. L. Papadopoulos, Y. Lubling, X. Xu, X. Lv, J. P. Hugnot, A. Tanay and G. Cavalli (2017). "Multiscale 3D Genome Rewiring during Mouse Neural Development." Cell 171(3): 557–572 e524.

Bouazoune, K. and R. E. Kingston (2012). "Chromatin remodeling by the CHD7 protein is impaired by mutations that cause human developmental disorders." Proc Natl Acad Sci U S A 109(47): 19238–19243.

Cao, X., S. L. Pfaff and F. H. Gage (2007). "A functional study of miR-124 in the developing neural tube." Genes Dev 21(5): 531–536.

Chang, L. H., S. Ghosh, A. Papale, J. M. Luppino, M. Miranda, V. Piras, J. Degrouard, J. Edouard, M. Poncelet, N. Lecouvreur, S. Bloyer, A. Leforestier, E. F. Joyce, D. Holcman and D. Noordermeer (2023). "Multi-feature clustering of CTCF binding creates robustness for loop extrusion blocking and Topologically Associating Domain boundaries." Nat Commun 14(1): 5615.

Chen, B., C. Ren, Z. Ouyang, J. Xu, K. Xu, Y. Li, H. Guo, X. Bai, M. Tian, X. Xu, Y. Wang, H. Li, X. Bo and H. Chen (2024). "Stratifying TAD boundaries pinpoints focal genomic regions of regulation, damage, and repair." Brief Bioinform 25(4).

Chen, W., N. Jongkamonwiwat, L. Abbas, S. J. Eshtan, S. L. Johnson, S. Kuhn, M. Milo, J. K. Thurlow, P. W. Andrews, W. Marcotti, H. D. Moore and M. N. Rivolta (2012). "Restoration of auditory evoked responses by human ES-cell-derived otic progenitors." Nature 490(7419): 278–282.

Coolen, M., S. Katz and L. Bally-Cuif (2013). "miR-9: a versatile regulator of neurogenesis." Front Cell Neurosci 7: 220.

Coolen, M., D. Thieffry, O. Drivenes, T. S. Becker and L. Bally-Cuif (2012). "miR-9 controls the timing of neurogenesis through the direct inhibition of antagonistic factors." Dev Cell 22(5): 1052–1064.

Corrales, C. E., L. Pan, H. Li, M. C. Liberman, S. Heller and A. S. Edge (2006). "Engraftment and differentiation of embryonic stem cell-derived neural progenitor cells in the cochlear nerve trunk: growth of processes into the organ of Corti." J Neurobiol 66(13): 1489–1500.

Dixon, J. R., S. Selvaraj, F. Yue, A. Kim, Y. Li, Y. Shen, M. Hu, J. S. Liu and B. Ren (2012). "Topological domains in mammalian genomes identified by analysis of chromatin interactions." Nature 485(7398): 376–380.

Ebert, M. S. and P. A. Sharp (2012). "Roles for microRNAs in conferring robustness to biological processes." Cell 149(3): 515–524.

Eling, N., M. D. Morgan and J. C. Marioni (2019). "Challenges in measuring and understanding biological noise." Nat Rev Genet 20(9): 536–548.

Elkan-Miller, T., I. Ulitsky, R. Hertzano, A. Rudnicki, A. A. Dror, D. R. Lenz, R. Elkon, M. Irmler, J. Beckers, R. Shamir and K. B. Avraham (2011). "Integration of transcriptomics, proteomics, and microRNA analyses reveals novel microRNA regulation of targets in the mammalian inner ear." PLoS One 6(4): e18195.

Engelen, E., U. Akinci, J. C. Bryne, J. Hou, C. Gontan, M. Moen, D. Szumska, C. Kockx, W. van Ijcken, D. H. Dekkers, J. Demmers, E. J. Rijkers, S. Bhattacharya, S. Philipsen, L. H. Pevny, F. G. Grosveld, R. J. Rottier, B. Lenhard and R. A. Poot (2011). "Sox2 cooperates with Chd7 to regulate genes that are mutated in human syndromes." Nat Genet 43(6): 607–611.

Friedman, L. M., A. A. Dror, E. Mor, T. Tenne, G. Toren, T. Satoh, D. J. Biesemeier, N. Shomron, D. M. Fekete, E. Hornstein and K. B. Avraham (2009). "MicroRNAs are essential for development and function of inner ear hair cells in vertebrates." Proc Natl Acad Sci U S A 106(19): 7915–7920.

Fujita, H., H. Aoki, I. Ajioka, M. Yamazaki, M. Abe, A. Oh-Nishi, K. Sakimura and I. Sugihara (2014). "Detailed expression pattern of aldolase C (Aldoc) in the cerebellum, retina and other areas of the CNS studied in Aldoc-Venus knock-in mice." PLoS One 9(1): e86679.

Gao, J., J. M. Skidmore, J. Cimerman, K. E. Ritter, J. Qiu, L. M. Q. Wilson, Y. Raphael, K. Y. Kwan and D. M. Martin (2024). "CHD7 and SOX2 act in a common gene regulatory network during mammalian semicircular canal and cochlear development." Proc Natl Acad Sci U S A 121(10): e2311720121.

Gerdes, J., U. Schwab, H. Lemke and H. Stein (1983). "Production of a mouse monoclonal antibody reactive with a human nuclear antigen associated with cell proliferation." Int J Cancer 31(1): 13–20.

Gu, Z. and D. Hubschmann (2023). "rGREAT: an R/bioconductor package for functional enrichment on genomic regions." Bioinformatics 39(1).

Gu, Z. and D. Hubschmann (2023). "simplifyEnrichment: A Bioconductor Package for Clustering and Visualizing Functional Enrichment Results." Genomics Proteomics Bioinformatics 21(1): 190–202.

Heinz, S., C. Benner, N. Spann, E. Bertolino, Y. C. Lin, P. Laslo, J. X. Cheng, C. Murre, H. Singh and C. K. Glass (2010). "Simple combinations of lineage-determining transcription factors prime cis-regulatory elements required for macrophage and B cell identities." Mol Cell 38(4): 576–589.

https://broadinstitute.github.io/picard/(2019). "Picard Toolkit." Broad Institute, GitHub repository.

Hurd, E. A., M. E. Adams, W. S. Layman, D. L. Swiderski, L. A. Beyer, K. E. Halsey, J. M. Benson, T. W. Gong, D. F. Dolan, Y. Raphael and D. M. Martin (2011). "Mature middle and inner ears express Chd7 and exhibit distinctive pathologies in a mouse model of CHARGE syndrome." Hear Res 282(1-2): 184–195.

Hurd, E. A., P. L. Capers, M. N. Blauwkamp, M. E. Adams, Y. Raphael, H. K. Poucher and D. M. Martin (2007). "Loss of Chd7 function in gene-trapped reporter mice is embryonic lethal and associated with severe defects in multiple developing tissues." Mamm Genome 18(2): 94–104.

Hurd, E. A., H. K. Poucher, K. Cheng, Y. Raphael and D. M. Martin (2010). "The ATP-dependent chromatin remodeling enzyme CHD7 regulates pro-neural gene expression and neurogenesis in the inner ear." Development 137(18): 3139–3150.

Ishihara, K., M. Oshimura and M. Nakao (2006). "CTCF-dependent chromatin insulator is linked to epigenetic remodeling." Mol Cell 23(5): 733–742.

Jacobs, S. A. and S. Khorasanizadeh (2002). "Structure of HP1 chromodomain bound to a lysine 9-methylated histone H3 tail." Science 295(5562): 2080–2083.

Jadali, A., Z. Song, A. S. Laureano, A. Toro-Ramos and K. Kwan (2016). "Initiating Differentiation in Immortalized Multipotent Otic Progenitor Cells." J Vis Exp(107).

Kabirova, E., A. Ryzhkova, V. Lukyanchikova, A. Khabarova, A. Korablev, T. Shnaider, M. Nuriddinov, P. Belokopytova, A. Smirnov, N. V. Khotskin, G. Kontsevaya, I. Serova and N. Battulin (2024). "TAD border deletion at the Kit locus causes tissue-specific ectopic activation of a neighboring gene." Nat Commun 15(1): 4521.

Kent, W. J., A. S. Zweig, G. Barber, A. S. Hinrichs and D. Karolchik (2010). "BigWig and BigBed: enabling browsing of large distributed datasets." Bioinformatics 26(17): 2204–2207.

Kim, D., B. Langmead and S. L. Salzberg (2015). "HISAT: a fast spliced aligner with low memory requirements." Nat Methods 12(4): 357–360.

Kim, J., E. Martinez, J. Qiu, J. Zhouli Ni and K. Y. Kwan (2024). "Chromatin remodeling protein CHD4 regulates axon guidance of spiral ganglion neurons in developing cochlea." bioRxiv.

Kim, K. L., G. J. Rahme, V. Y. Goel, C. A. El Farran, A. S. Hansen and B. E. Bernstein (2024). "Dissection of a CTCF topological boundary uncovers principles of enhancer-oncogene regulation." Mol Cell 84(7): 1365–1376 e1367.

Kim, T. H., Z. K. Abdullaev, A. D. Smith, K. A. Ching, D. I. Loukinov, R. D. Green, M. Q. Zhang, V. V. Lobanenkov and B. Ren (2007). "Analysis of the vertebrate insulator protein CTCF-binding sites in the human genome." Cell 128(6): 1231–1245.

Kloosterman, W. P., E. Wienholds, E. de Bruijn, S. Kauppinen and R. H. Plasterk (2006). "In situ detection of miRNAs in animal embryos using LNA-modified oligonucleotide probes." Nat Methods 3(1): 27–29.

Kouzarides, T. (2007). "Chromatin modifications and their function." Cell 128(4): 693–705.

Kubo, N., H. Ishii, X. Xiong, S. Bianco, F. Meitinger, R. Hu, J. D. Hocker, M. Conte, D. Gorkin, M. Yu, B. Li, J. R. Dixon, M. Hu, M. Nicodemi, H. Zhao and B. Ren (2021). "Promoter-proximal CTCF binding promotes distal enhancer-dependent gene activation." Nat Struct Mol Biol 28(2): 152–161.

Kujawa, S. G. and M. C. Liberman (2006). "Acceleration of age-related hearing loss by early noise exposure: evidence of a misspent youth." J Neurosci 26(7): 2115–2123.

Kujawa, S. G. and M. C. Liberman (2009). "Adding insult to injury: cochlear nerve degeneration after "temporary" noise-induced hearing loss." J Neurosci 29(45): 14077–14085.

Kutner, R. H., X. Y. Zhang and J. Reiser (2009). "Production, concentration and titration of pseudotyped HIV-1-based lentiviral vectors." Nat Protoc 4(4): 495–505.

Kwan, K. Y., J. Shen and D. P. Corey (2015). "C-MYC transcriptionally amplifies SOX2 target genes to regulate self-renewal in multipotent otic progenitor cells." Stem Cell Reports 4(1): 47–60.

Lagos-Quintana, M., R. Rauhut, A. Yalcin, J. Meyer, W. Lendeckel and T. Tuschl (2002). "Identification of tissue-specific microRNAs from mouse." Curr Biol 12(9): 735–739.

Langmead, B. and S. L. Salzberg (2012). "Fast gapped-read alignment with Bowtie 2." Nat Methods 9(4): 357–359.

Li, B., M. Carey and J. L. Workman (2007). "The role of chromatin during transcription." Cell 128(4): 707–719.

Li, C., X. Li, Z. Bi, K. Sugino, G. Wang, T. Zhu and Z. Liu (2020). "Comprehensive transcriptome analysis of cochlear spiral ganglion neurons at multiple ages." Elife 9.

Li, H., B. Handsaker, A. Wysoker, T. Fennell, J. Ruan, N. Homer, G. Marth, G. Abecasis, R. Durbin and S. Genome Project Data Processing (2009). "The Sequence Alignment/Map format and SAMtools." Bioinformatics 25(16): 2078–2079.

Li, W., Y. Xiong, C. Shang, K. Y. Twu, C. T. Hang, J. Yang, P. Han, C. Y. Lin, C. J. Lin, F. C. Tsai, K. Stankunas, T. Meyer, D. Bernstein, M. Pan and C. P. Chang (2013). "Brg1 governs distinct pathways to direct multiple aspects of mammalian neural crest cell development." Proc Natl Acad Sci U S A 110(5): 1738–1743.

Love, M. I., W. Huber and S. Anders (2014). "Moderated estimation of fold change and dispersion for RNA-seq data with DESeq2." Genome Biol 15(12): 550.

Lu, C. C., J. M. Appler, E. A. Houseman and L. V. Goodrich (2011). "Developmental profiling of spiral ganglion neurons reveals insights into auditory circuit assembly." J Neurosci 31(30): 10903–10918.

Lupianez, D. G., K. Kraft, V. Heinrich, P. Krawitz, F. Brancati, E. Klopocki, D. Horn, H. Kayserili, J. M. Opitz, R. Laxova, F. Santos-Simarro, B. Gilbert-Dussardier, L. Wittler, M. Borschiwer, S. A. Haas, M. Osterwalder, M. Franke, B. Timmermann, J. Hecht, M. Spielmann, A. Visel and S. Mundlos (2015). "Disruptions of topological chromatin domains cause pathogenic rewiring of gene-enhancer interactions." Cell 161(5): 1012–1025.

Manning, B. J. and T. Yusufzai (2017). "The ATP-dependent chromatin remodeling enzymes CHD6, CHD7, and CHD8 exhibit distinct nucleosome binding and remodeling activities." J Biol Chem 292(28): 11927–11936.

Meers, M. P., D. Tenenbaum and S. Henikoff (2019). "Peak calling by Sparse Enrichment Analysis for CUT&RUN chromatin profiling." Epigenetics Chromatin 12(1): 42.

Mukherji, S., M. S. Ebert, G. X. Zheng, J. S. Tsang, P. A. Sharp and A. van Oudenaarden (2011). "MicroRNAs can generate thresholds in target gene expression." Nat Genet 43(9): 854–859.

Nanni, L., S. Ceri and C. Logie (2020). "Spatial patterns of CTCF sites define the anatomy of TADs and their boundaries." Genome Biol 21(1): 197.

Nielsen, P. R., D. Nietlispach, H. R. Mott, J. Callaghan, A. Bannister, T. Kouzarides, A. G. Murzin, N. V. Murzina and E. D. Laue (2002). "Structure of the HP1 chromodomain bound to histone H3 methylated at lysine 9." Nature 416(6876): 103–107.

Nishimura, K., T. Nakagawa, T. Sakamoto and J. Ito (2012). "Fates of murine pluripotent stem cell-derived neural progenitors following transplantation into mouse cochleae." Cell Transplant 21(4): 763–771.

Nitiss, J. L. (1998). "Investigating the biological functions of DNA topoisomerases in eukaryotic cells." Biochim Biophys Acta 1400(1-3): 63–81.

Nora, E. P., A. Goloborodko, A. L. Valton, J. H. Gibcus, A. Uebersohn, N. Abdennur, J. Dekker, L. A. Mirny and B. G. Bruneau (2017). "Targeted Degradation of CTCF Decouples Local Insulation of Chromosome Domains from Genomic Compartmentalization." Cell 169(5): 930–944 e922.

Petitpre, C., L. Faure, P. Uhl, P. Fontanet, I. Filova, G. Pavlinkova, I. Adameyko, S. Hadjab and F. Lallemend (2022). "Single-cell RNA-sequencing analysis of the developing mouse inner ear identifies molecular logic of auditory neuron diversification." Nat Commun 13(1): 3878.

Quinlan, A. R. and I. M. Hall (2010). "BEDTools: a flexible suite of utilities for comparing genomic features." Bioinformatics 26(6): 841–842.

Ramirez, F., F. Dundar, S. Diehl, B. A. Gruning and T. Manke (2014). "deepTools: a flexible platform for exploring deep-sequencing data." Nucleic Acids Res 42(Web Server issue): W187–191.

Rao, S. S., M. H. Huntley, N. C. Durand, E. K. Stamenova, I. D. Bochkov, J. T. Robinson, A. L. Sanborn, I. Machol, A. D. Omer, E. S. Lander and E. L. Aiden (2014). "A 3D map of the human genome at kilobase resolution reveals principles of chromatin looping." Cell 159(7): 1665–1680.

Robinson, J. T., H. Thorvaldsdottir, W. Winckler, M. Guttman, E. S. Lander, G. Getz and J. P. Mesirov (2011). "Integrative genomics viewer." Nat Biotechnol 29(1): 24–26.

Rudnicki, A. and K. B. Avraham (2012). "microRNAs: the art of silencing in the ear." EMBO Mol Med 4(9): 849–859.

Sanders, T. R. and M. W. Kelley (2022). "Specification of neuronal subtypes in the spiral ganglion begins prior to birth in the mouse." Proc Natl Acad Sci U S A 119(48): e2203935119.

Sanosaka, T., H. Okuno, N. Mizota, T. Andoh-Noda, M. Sato, R. Tomooka, S. Banno, J. Kohyama and H. Okano (2022). "Chromatin remodeler CHD7 targets active enhancer region to regulate cell type-specific gene expression in human neural crest cells." Sci Rep 12(1): 22648.

Schmiedel, J. M., S. L. Klemm, Y. Zheng, A. Sahay, N. Bluthgen, D. S. Marks and A. van Oudenaarden (2015). "Gene expression. MicroRNA control of protein expression noise." Science 348(6230): 128–132.

Schnetz, M. P., C. F. Bartels, K. Shastri, D. Balasubramanian, G. E. Zentner, R. Balaji, X. Zhang, L. Song, Z. Wang, T. Laframboise, G. E. Crawford and P. C. Scacheri (2009). "Genomic distribution of CHD7 on chromatin tracks H3K4 methylation patterns." Genome Res 19(4): 590–601.

Selbach, M., B. Schwanhausser, N. Thierfelder, Z. Fang, R. Khanin and N. Rajewsky (2008). "Widespread changes in protein synthesis induced by microRNAs." Nature 455(7209): 58–63.

Shi, F. and A. S. Edge (2013). "Prospects for replacement of auditory neurons by stem cells." Hear Res 297: 106–112.

Shibata, M., H. Nakao, H. Kiyonari, T. Abe and S. Aizawa (2011). "MicroRNA-9 regulates neurogenesis in mouse telencephalon by targeting multiple transcription factors." J Neurosci 31(9): 3407–3422.

Soukup, G. A., B. Fritzsch, M. L. Pierce, M. D. Weston, I. Jahan, M. T. McManus and B. D. Harfe (2009). "Residual microRNA expression dictates the extent of inner ear development in conditional Dicer knockout mice." Dev Biol 328(2): 328–341.

Strahl, B. D. and C. D. Allis (2000). "The language of covalent histone modifications." Nature 403(6765): 41-45.

Tarjan, D. R., W. A. Flavahan and B. E. Bernstein (2019). "Epigenome editing strategies for the functional annotation of CTCF insulators." Nat Commun 10(1): 4258.

Thelin, J. W. and J. C. Fussner (2005). "Factors related to the development of communication in CHARGE syndrome." Am J Med Genet A 133A(3): 282-290.

Thelin, J. W., J. A. Mitchell, M. A. Hefner and S. L. Davenport (1986). "CHARGE syndrome. Part II. Hearing loss." Int J Pediatr Otorhinolaryngol 12(2): 145–163.

Vissers, L. E., C. M. van Ravenswaaij, R. Admiraal, J. A. Hurst, B. B. de Vries, I. M. Janssen, W. A. van der Vliet, E. H. Huys, P. J. de Jong, B. C. Hamel, E. F. Schoenmakers, H. G. Brunner, J. A. Veltman and A. G. van Kessel (2004). "Mutations in a new member of the chromodomain gene family cause CHARGE syndrome." Nat Genet 36(9): 955–957.

Weston, M. D., M. L. Pierce, S. Rocha-Sanchez, K. W. Beisel and G. A. Soukup (2006). "MicroRNA gene expression in the mouse inner ear." Brain Res 1111(1): 95–104.

Xiao, J. Y., A. Hafner and A. N. Boettiger (2021). "How subtle changes in 3D structure can create large changes in transcription." Elife 10.

Yuva-Aydemir, Y., A. Simkin, E. Gascon and F. B. Gao (2011). "MicroRNA-9: functional evolution of a conserved small regulatory RNA." RNA Biol 8(4): 557–564.

Zaghi, M., F. Banfi, L. Massimino, M. Volpin, E. Bellini, S. Brusco, I. Merelli, C. Barone, M. Bruni, L. Bossini, L. A. Lamparelli, L. Pintado, D. D’Aliberti, S. Spinelli, L. Mologni, G. Colasante, F. Ungaro, J. M. Cioni, E. Azzoni, R. Piazza, E. Montini, V. Broccoli and A. Sessa (2023). "Balanced SET levels favor the correct enhancer repertoire during cell fate acquisition." Nat Commun 14(1): 3212.

Zentner, G. E., W. S. Layman, D. M. Martin and P. C. Scacheri (2010). "Molecular and phenotypic aspects of CHD7 mutation in CHARGE syndrome." Am J Med Genet A 152A(3): 674–686.

